# The iron-responsive genome of the chiton *Acanthopleura granulata*

**DOI:** 10.1101/2020.05.19.102897

**Authors:** Rebecca M. Varney, Daniel I. Speiser, Carmel McDougall, Bernard M. Degnan, Kevin M. Kocot

**Affiliations:** The University of Alabama, Department of Biological Sciences; University of South Carolina, Department of Biological Sciences; Griffith University, Australian Rivers Institute; University of Queensland, School of Biological Sciences; The University of Alabama Museum of Natural History

**Keywords:** biomineralization, biomaterials, iron, ferritin, Iron response element (IRE), mollusc

## Abstract

Molluscs biomineralize structures that vary in composition, form, and function, prompting questions about the genetic mechanisms responsible for their production and the evolution of these mechanisms. Chitons (Mollusca, Polyplacophora) are a promising system for studies of biomineralization because they build a range of calcified structures including shell plates and spine- or scale-like sclerites. Chitons also harden the calcified teeth of their rasp-like radula with a coat of iron (as magnetite). Here we present the genome of the West Indian fuzzy chiton *Acanthopleura granulata*, the first from any aculiferan mollusc. The *A. granulata* genome contains homologs of many biomineralization genes identified previously in conchiferan molluscs. We expected chitons to lack genes previously identified from pathways conchiferans use to make biominerals like calcite and nacre because chitons do not use these materials in their shells. Surprisingly, the *A. granulata* genome has homologs of many of these genes, suggesting that the ancestral mollusc had a more diverse biomineralization toolkit than expected. The *A. granulata* genome has features that may be specialized for iron biomineralization, including a higher proportion of genes regulated directly by iron than other molluscs. *A. granulata* also produces two isoforms of soma-like ferritin: one is regulated by iron and similar in sequence to the soma-like ferritins of other molluscs, and the other is constitutively translated and is not found in other molluscs. The *A. granulata* genome is a resource for future studies of molluscan evolution and biomineralization.

**SIGNIFICANCE STATEMENT:** Chitons are molluscs that make shell plates, spine- or scale-like sclerites, and iron-coated teeth. Currently, all molluscs with sequenced genomes lie within one major clade (Conchifera). Sequencing the genome of a representative from the other major clade (Aculifera) helps us learn about the origins and evolution of molluscan traits. The genome of the West Indian Fuzzy Chiton, *Acanthopleura granulata*, reveals chitons have homologs of many genes other molluscs use to make shells, suggesting all molluscs share some shell-making pathways. The genome of *A. granulata* has more genes that may be regulated directly by iron than other molluscs, and chitons produce a unique isoform of a major iron-transport protein (ferritin), suggesting that chitons have genomic specializations that contribute to their production of iron-coated teeth.

## INTRODUCTION

Animals construct hardened structures by combining organic and inorganic components, a process termed biomineralization. To do so, they secrete proteins that initiate and guide the crystallization of inorganic molecules. Animals also incorporate proteins into biomineralized structures, enhancing their strength and flexibility (Cölfen 2010). Molluscs have long been models for studying the genetic mechanisms associated with biomineralization because they craft a wide range of materials into shells, spines, scales, and teeth (McDougall & Degnan 2018). The ability of molluscs to produce diverse biomineralized structures likely contributes to their remarkable morphological and ecological diversity.

Chitons (Polyplacophora, Figure 1A) are a promising model for investigating mechanisms of biomineralization because they build diverse mineralized structures distinct from those of other molluscs. The shells of all molluscs are composed of calcium carbonate (CaCO_3_), commonly in its crystal forms aragonite or calcite. Most molluscs build shells with alternating layers of aragonite and calcite, and many add an innermost layer of brick-like aragonite discs known as nacre. In contrast, chitons construct eight interlocking shell plates (Figure 1B) exclusively from aragonite and do not produce nacre. Also unlike other molluscs, chitons embed a network of sensory structures, termed aesthetes, into their shell plates. In some species, the aesthete network includes eyes with image-forming lenses made of aragonite (Speiser et al. 2011; Li et al. 2015); Figure 1C). To protect the soft girdle tissue surrounding their shell plates, chitons produce scale- or spine-like sclerites, which are also made of aragonite (Schwabe 2010; Sigwart et al. 2014; Checa et al. 2017).

**Figure 1:**
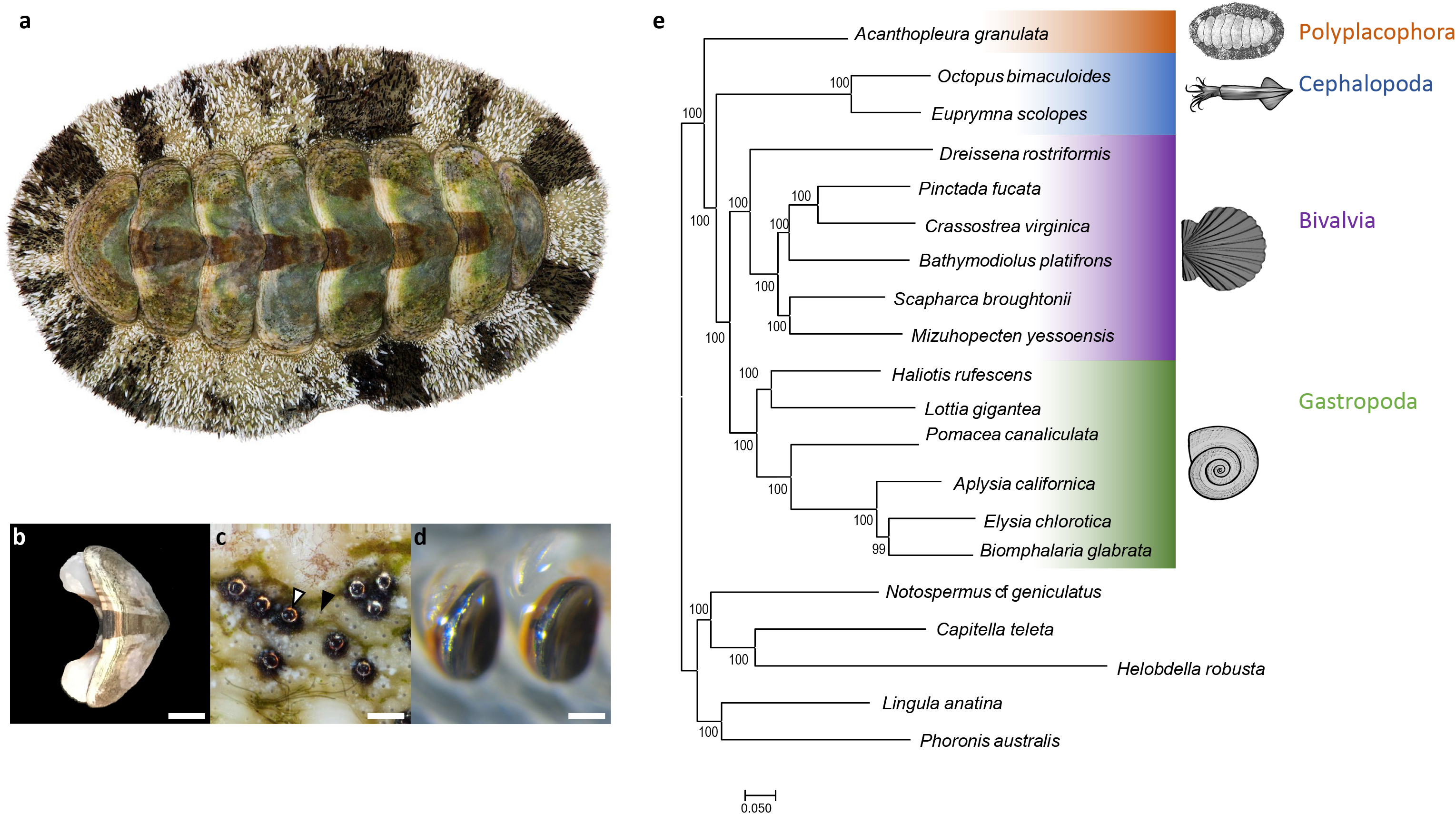
(a) The West Indian Fuzzy Chiton Acanthopleura granulata. Photograph by David Liittschwager. (b) A single shell plate from A. granulata. Scale bar indicates 5mm. (c) The eyes (white arrow) and aesthetes (black arrow) of A. granulata. Scale bar indicates 200 μM. Photograph by David Liittschwager. (d) Teeth from the anterior-most region of the radula of A. granulata. The larger teeth, used for feeding, are mineralized with iron oxide (orange) and capped with magnetite (black). Scale bar indicates 300 μM (e) A genome-based phylogeny of Mollusca showing chitons as sister to all other molluscs with available genomes.

Chitons biomineralize their teeth (Figure 1D) from a unique combination of materials. Most molluscs have a feeding organ, the radula, that bears rows of teeth built from chitin and, in many species, hardened with minerals such as calcium carbonate or silica. Chitons instead harden the cores of their teeth with calcium phosphate (as apatite), and then reinforce their cutting edges with iron (as magnetite) (Lowenstam 1962). These iron coatings allow chitons to scrape algae from rocks without rapidly dulling or damaging their teeth. Chitons produce new teeth throughout their lives, making new rows within days (Shaw et al. 2002; Joester & Brooker 2016). To make new teeth, chitons continuously sequester and transport high concentrations of iron (Kim et al. 1989; Shaw et al. 2002, 2010). Continuous iron biomineralization may thus present a physiological challenge to chitons because free iron causes oxidative stress (Dixon & Stockwell 2014).

To date, most investigations of biomineralization in molluscs have focused on species from the classes Bivalvia and Gastropoda. These, together with Monoplacophora, Cephalopoda, and Scaphopoda make up the clade Conchifera. The sister clade to Conchifera is Aculifera, which is made up of Polyplacophora and Aplacophora. Conchifera and Aculifera diverged approximately 550mya (Vinther et al. 2012; Kocot et al. 2020). To make robust predictions about molluscan evolution, reconstructions of ancestral character states must include information from both conchiferans and aculiferans (Sigwart & Sutton 2007; Kocot et al. 2011; Smith et al. 2011; Vinther et al. 2012). Despite increasing numbers of sequenced molluscan genomes (e.g., Takeuchi et al. 2012; Zhang et al. 2012; Simakov et al. 2013; Albertin et al. 2015; Gómez-Chiarri et al. 2015; Modica et al. 2015; Kenny et al. 2015; Barghi et al. 2016; Davison et al. 2016; Murgarella et al. 2016; Adema et al. 2017; Du et al. 08 01, 2017; Nam et al. 05 01, 2017; Schell et al. 2017; Sun et al. 2017; Wang et al. 2017; Calcino et al. 2018; Gerdol et al. 05 2018; Li et al. 04 01, 2018; Liu et al. 2018; Renaut et al. 07 01, 2018; Belcaid et al. 2019; Cai et al. 2019; Kijas et al. 2019; Masonbrink et al. 2019; McCartney et al. 2019; Zarrella et al. 2019; Sun et al. 2020), genomic resources for aculiferans remain unavailable. To advance the study of molluscan evolution and to better understand the genetic mechanisms of biomineralization, we sequenced the genome of the West Indian fuzzy chiton *Acanthopleura granulata*. Exploring the *A. granulata* genome allowed us to: 1) identify genes chitons may use to build their shell plates, sclerites, and teeth; 2) seek genomic signatures associated with the biomineralization of iron and the mitigation of iron-induced oxidative stress; and 3) better understand the origin and evolution of biomineralization in molluscs.

## RESULTS AND DISCUSSION

We sequenced the genome of a single specimen of *A. granulata*. We combined reads from one lane of Illumina HiSeq X paired-end sequencing (124 Gb of 2 × 150 bp reads,~204X coverage) with reads from four Oxford Nanopore flowcells run on the GridION platform (22.87 Gb, 37X coverage). Using the hybrid assembler MaSuRCA and optical mapping, we produced a haploid genome assembly for *A. granulata* that is 606.9 Mbp, slightly smaller than the 743 Mbp haploid genome size estimated by flow cytometry (Roebuck 2017). The assembled *A. granulata* genome consists of 87 scaffolds ranging in size from 50.9 Mb to 0.05 Mb, plus a single mitochondrial genome of 15,665 bp. Several of these scaffolds are similar in length to intact chromosomes from other molluscs (Sun et al. 2017, 2020; Bai et al. 2019). To verify completeness of the assembly, we mapped genomic short-read data to the genome; 85.31% of reads mapped perfectly, so we are confident the assembly encompasses a majority of sequencing data. The *A. granulata* genome has an N50 value of 23.9 Mbp and a BUSCO completeness score of 97.4%, making it more contiguous and complete than most currently available molluscan genomes (Supplementary Figure 1; Supplementary Table 1; visualized in Supplementary Figure 2).

We generated gene models for *A. granulata* by 1) sequencing transcriptomes from eight different tissues from the same specimen used for genome sequencing, 2) combining these transcriptomes into a single assembly and aligning the combined transcriptome to the genome, and 3) training *de novo* gene predictors using both our combined transcriptome and protein sequences predicted from the transcriptomes of other aculiferans. Following these steps, we produced a set of 81,691 gene models that is 96.9% complete according to a BUSCO transcriptomic analysis. This score is similar to the completeness score of the *A. granulata* genome, so it is likely this set of gene models missed few genes, if any, in the genome assembly. However, of the BUSCO genes expected to be single-copy in all animals, 17.2% were represented by multiple gene models. Using Markov clustering to eliminate redundant isoforms, we generated a reduced set of 20,470 gene models that is 94.7% complete. In this smaller set of gene models, only 0.5% of the BUSCO genes have multiple copies, supporting Markov clustering as an effective method for reducing the redundancy of gene models. To characterize proteins based on shared functional domains and sequence similarity, we analyzed the set of 20,470 gene models with InterProScan. We identified at least one GO term for 12,301 genes and a Pfam match for 15,710 genes. We also conducted a KEGG analysis and identified 7,341 proteins that could be assigned to putative molecular pathways.

To provide a robust dataset for phylogenetic analysis and gene family evolution analyses, we identified homologous genes shared between *A. granulata* and other molluscs. We used the larger set of gene models from *A. granulata* to ensure a more complete input dataset, knowing that any duplicate gene models for the same locus would cluster within the same orthologous group. We compared gene models from the *A. granulata* genome to those from the genomes of nineteen other lophotrochozoans, including fourteen molluscs, two annelids, one brachiopod, one phoronid, and one nemertean. This resulted in 59,276 groups of homologous sequences including 3,379 found in all 20 genomes.

We used a tree-based approach to identify orthologous genes shared among all 20 taxa and reconstructed molluscan phylogeny using the 2,593 orthologs present in at least 17 of the 20 genomes we searched. This dataset totaled 950,322 amino acid positions with 16.2% missing data. We recovered *A. granulata* as the sister taxon of all other molluscs with sequenced genomes (Figure 1F). We conducted an additional phylogenetic analysis that included more taxa by using transcriptomes in addition to genomes and recovered *Acanthopleura* within the family Chitonidae in the order Chitonida, consistent with recent phylogenetic studies of chitons based on fewer loci (Irisarri et al. 2020); Supplementary Figure 3).

### The *A. granulata* genome differs from conchiferan genomes in content and organization

The *A. granulata* genome has a heterozygosity of 0.653%, making it one of the least heterozygous molluscan genomes sequenced to date (Supplementary Figure 4). High heterozygosity is often attributed to high rates of gene flow associated with broadcast spawning and far-dispersing larvae (Solé-Cava & Thorpe 1991), and it is frequently noted as an obstacle to genome assembly in molluscs (Zhang et al. 2012; Wang et al. 2017; Powell et al. 2018; Thai et al. 2019). We expected the genome of *A. granulata* to have high heterozygosity because this species is a broadcast spawner with a wide geographic range (Glynn 1970). To compare heterozygosity across molluscs, we selected a set of high-quality molluscan genomes for which short-read data are available (Supplementary Table 1). Using k-mer based analysis, we found the highest heterozygosity among the seven genomes we analyzed was 3.15% in the blood clam *S. broughtonii*, and the other genomes had heterozygosities between those of *A. granulata* and *S. broughtonii*. Our findings indicate that heterozygosity may be influenced by more than an animal’s reproductive mode, larval type, and geographic range (Supplementary Table 2), and that molluscan genomes should not be assumed to have high heterozygosity.

The *A. granulata* genome is arranged differently than other molluscan genomes and has fewer repetitive elements. Compared to a non-molluscan lophotrochozoan, *Lingula anatina* (a brachiopod), *A. granulata* has more repetitive elements of certain types in its genome. Conversely, *A. granulata* has fewer repetitive elements in its genome than any conchiferan mollusc (Supplementary Table 2). This suggests multiple proliferations of repetitive elements during molluscan evolution. Repetitive elements contribute to structural changes in genomes by providing breakpoints that increase the likelihood of chromosomal rearrangements (Weckselblatt & Rudd 2015). Consistent with this prediction, synteny is lower between *A. granulata* and all conchiferan molluscs we examined than it is between any two of these conchiferans, and the genomes of conchiferans and *A. granulata* have little synteny with the genome of *L. anatina* (Supplementary Figure 5). A recent study found greater conservation of bilaterian ancestral linkage groups (ALGs) between a scallop and non-molluscs than between other bivalves and non-molluscs, suggesting substantial genomic rearrangements occurred within bivalves (Wang et al. 2017). Molluscan genomes appear to rearrange frequently across evolutionary time, and perhaps rearrange more frequently in conchiferans due to the proliferation of repetitive elements. However, limited synteny between *A. granulata* and the scallop genome suggests that the chiton genome has also undergone significant rearrangement relative to the hypothetical ancestral bilaterian genome organization.

The Hox cluster is a widely conserved set of regulatory genes that contribute to the patterning of the anterior-posterior axes in bilaterian animals. In lophotrochozoans, the genes are typically collinear, beginning with *Hox1* and ending with *Post1*. Although several gastropods and bivalves possess intact Hox clusters, this cluster is dispersed in some bivalves and cephalopods (Albertin et al. 2015; Barucca et al. 2016; Belcaid et al. 2019; Wang et al. 2017). The Hox cluster of *A. granulata* lacks *Post1*, but is otherwise intact and collinear (Figure 2). Given current understanding of molluscan phylogeny, the order of Hox genes shared between *A. granulata* and most conchiferans likely represents the ancestral order of Hox genes in molluscs (Wanninger & Wollesen 2019); Figure 2). *Post1* is also absent in two species of chitons from the suborder Acanthochitonina, the sister clade to Chitonina, to which *A. granulata* belongs (Huan et al. 2019; Wanninger & Wollesen 2019). However, *Post1* is present in aplacophorans (Iijima et al. 2006), suggesting it was lost in chitons. In conchiferan molluscs, *Post1* helps specify the posterior of an animal during development and helps pattern shell formation (Lee et al. 2003; Fröbius et al. 2008; Schiemann et al. 2017; Huan et al. 2019). In the absence of *Post1, A. granulata* and other chitons must use other transcription factors to help pattern their body axes and biomineralized structures.

**Figure 2:**
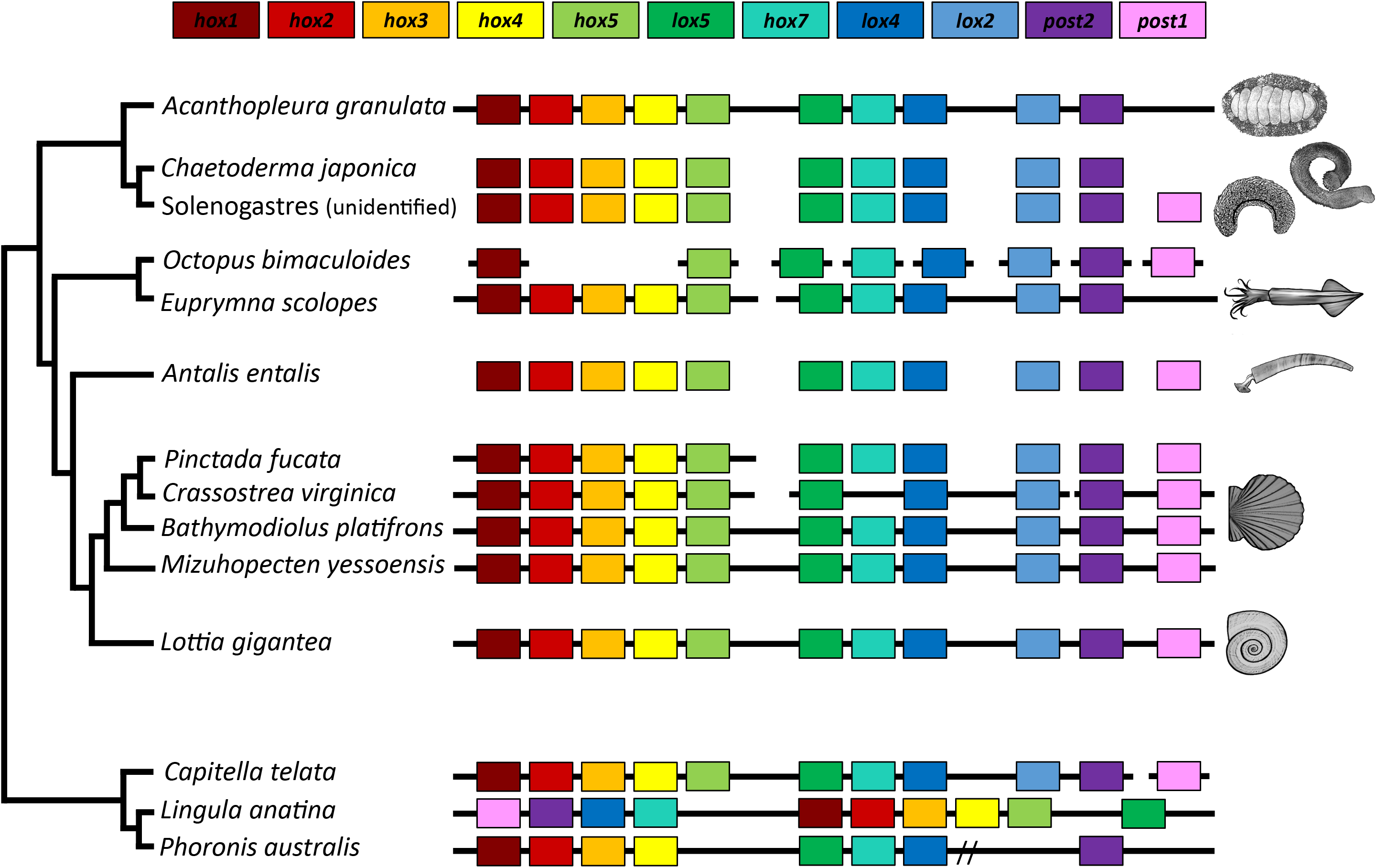
Synteny of Hox genes between A. granulata and other taxa. The presence of a gene is indicated by a box of the corresponding color. Continuous black lines indicate that the species has an available genome and Hox genes were located on a contiguous scaffold. Broken black lines indicate that gene(s) are located on multiple genomic scaffolds. A double slash indicates genes are located on a single contiguous scaffold but separated by greater distances than those in most other taxa.

### *Acanthopleura granulata* shares many biomineralization genes with conchiferan molluscs

We expected chitons to lack many genes previously identified in molluscan biomineralization pathways because their shell plates and sclerites lack both calcite and nacre. We were surprised to find homologs in the *A. granulata* genome of many biomineralization genes known from conchiferans (Supplementary Table 4). For example, we found an ortholog in *A. granulata* for *Pif*. In pterid bivalves, the *Pif* mRNA encodes a protein that is cleaved into two peptides, PIF97 and PIF80 (Suzuki et al. 2009). These peptides have different roles in biomineralization: PIF80 binds nacre and aids in nacre formation (Suzuki et al. 2009), whereas PIF97 binds to chitin and guides the growth of calcium carbonate crystals (Suzuki et al. 2013). A recent study in the gastropod *Lymnaea stagnalis* identified a PIF-like protein that does not promote nacre production, but is expressed in tissues that suggest this PIF protein still plays a role in shell biomineralization (Ishikawa et al. 2020). We found that *A. granulata* possesses a *Pif* homolog, but appears to only produce PIF97 rather than two separate peptides. The expression of *Pif* mRNA was highest in girdle tissue in *A. granulata* and lowest in the radula, suggesting that PIF peptides may play a role in sclerite formation in chitons (Supplementary Table 4). We hypothesize that the last common ancestor of extant molluscs used PIF97 to help build mineralized structures, and that production of PIF80 is novel to bivalves.

The ancestral mollusc likely produced mineralized structures, but whether the ancestral mollusc had a single shell, multiple shell plates, or sclerites remains a matter of debate (Scherholz et al. 2013; Vinther et al. 2017; Giribet & Edgecombe 2020; Kocot 01/2013). Molluscs form mineralized structures by making extracellular matrices from organic components such as polysaccharides and proteins, and then hardening them with minerals (Furuhashi et al. 2009). Similarities between the extracellular matrices of different biomineralized structures suggest these structures share developmental mechanisms. The *A. granulata* genome includes genes known from conchiferan molluscs to be associated with extracellular matrices. Chitin is a major component of the extracellular matrices of all molluscan shells and radulae, and the *A. granulata* genome contains genes for chitin synthase, chitinase, and chitin-binding proteins. We also found homologs of lustrin and dermatopontin, two proteins expressed in the extracellular matrices of conchiferans that increase the elasticity and flexibility of their shells (Gaume et al. 2014); Supplementary Table 5).

Silk-like structural proteins are components of many biological materials, including shells (Eisoldt et al. 2011; McDougall et al. 2016; Xu et al. 2016), and several *A. granulata* genes are similar to genes known to code for silk-like proteins. These proteins are “silk-like” because they contain highly repetitive sequences of amino acids that fold into secondary structures (commonly β-pleated sheets) that impart flexibility, a phenomenon first documented in spider silk (Lewis 2006; Eisoldt et al. 2011). Silk-like domains can facilitate the precipitation and crystallization of minerals that help form structures such as bones and shells (Xu et al. 2016). We found 31 genes that code for proteins with silk-like domains in the *A. granulata* genome, 23 of which have high sequence similarity to characterized molluscan biomineralization genes (Supplementary Table 6). We found 27 of these 31 genes code for proteins with signal peptides, indicating they may be secreted as part of the extracellular matrix during biomineralization (Supplementary Table 6). Among these genes, we found three collagens, one chitinase, and one carbonic anhydrase, all possible contributors to shell formation and repair (Patel 2004); Supplementary Table 6). Several of the genes encoding proteins with silk-like domains are highly expressed in the girdle of *A. granulata*, suggesting a role in the mineralization of sclerites (Supplementary Figure 6).

### *A. granulata* has more genes with iron response elements (IREs) than other molluscs

Chitons have more iron in their hemolymph than any other animal studied to date (Kim et al. 1988). Iron presents physiological challenges to animals because iron can cause oxidative stress. We hypothesize that the ability of chitons to biomineralize iron requires them to respond quickly to changes in concentration of this potentially toxic metal. To assess the iron-responsiveness of the *A. granulata* genome, we searched it for iron response elements (IREs), three-dimensional hairpin structures that form in either the 3’ or 5’ untranslated regions (UTRs) of mRNA molecules and control translation via binding by iron regulatory protein (IRP; Supplementary Figure 7). We also examined IREs in several high-quality molluscan genomes that include UTRs as part of their available annotation data. All of the molluscan genomes we examined had similar proportions of 3’ to 5’ IREs (Figure 3A). Despite having the fewest gene models, the genome of *A. granulata* has more IREs than the genomes of any other mollusc we examined. We predicted 271 IREs in the *A. granulata* genome, compared with an average of 119 IREs across other molluscan genomes (Supplementary Table 7). The highest number of predicted IREs in a conchiferan came from the genome of the blood clam *Scapharca broughtonii*, which had 201. The blood clam is so named because it is one of relatively few molluscs that produces hemoglobin for use as a respiratory pigment (Kawamoto 1928; Manwell 1963; Read 1966; Collett & O’Gower 1972; Bai et al. 2019). We expect *A. granulata* and *S. broughtonii* have more IREs in their genomes than other molluscs because they must absorb and transport larger amounts of iron to produce iron-coated teeth and hemoglobin, respectively.

**Figure 3:**
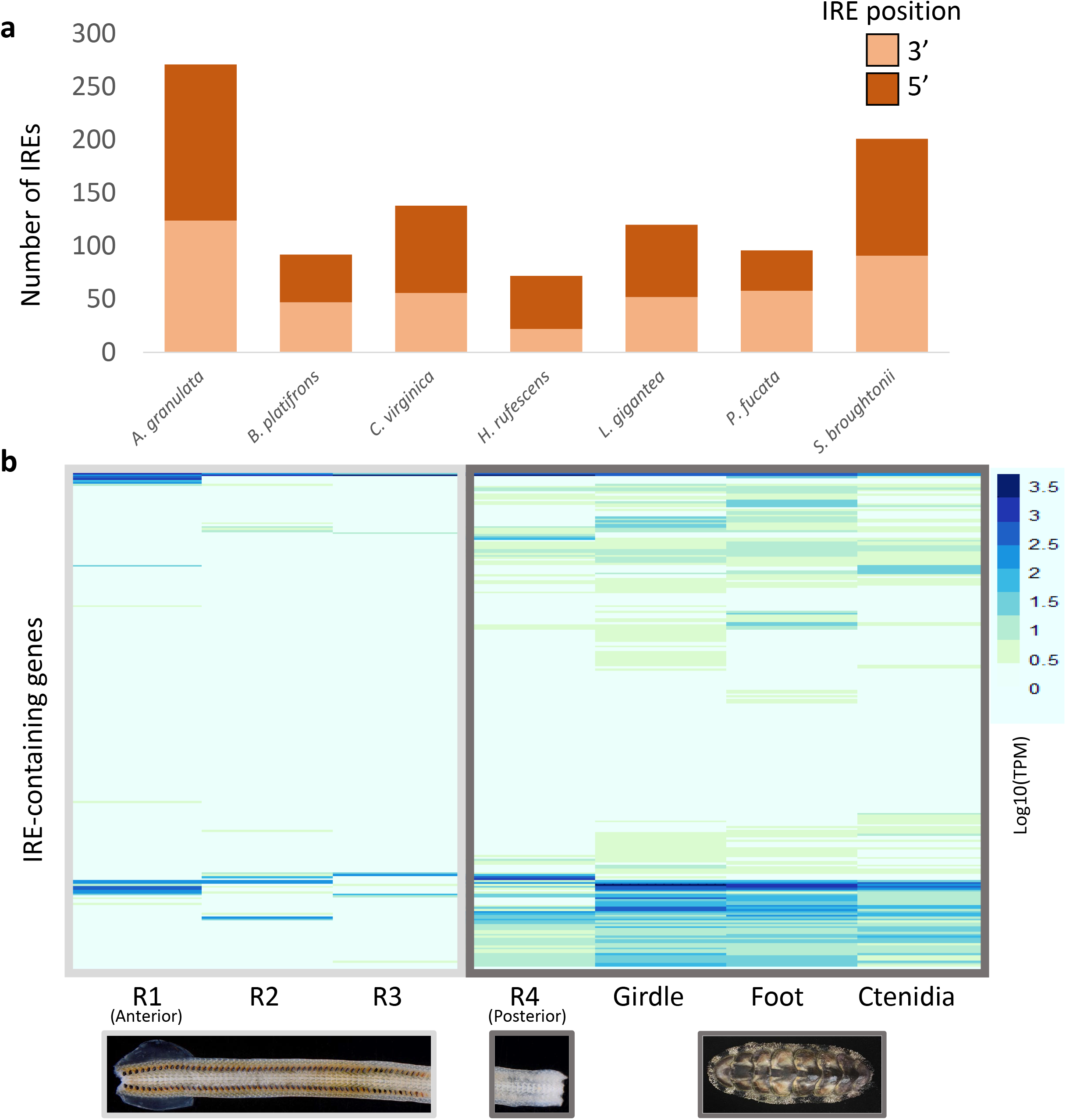
Iron response elements (IREs) in the A. granulata genome. (a) The number of IREs in several molluscan gene model sets, and relative proportions of 5’ and 3’ IREs. A. granulata has more IREs than all other molluscs examined, but the relative proportion of 5’ and 3’ IREs appears consistent across molluscan genomes. (b) The relative expression [log10(TPM)] of transcripts containing IREs in the different tissues of A. granulata. The radula is divided into four developmentally distinct regions: R1,the most anterior region, contains teeth used for feeding; R2 contains teeth that are developed but are not yet used for feeding; R3 contains developing teeth that contain iron oxide; and R4, the most posterior region, contains developing teeth that have yet to be coated with iron. We found lower expression of most IRE-containing genes in the anterior regions of the radula. (LogTPM used to allow both data ranges to appear legibly on the same graph)

We next asked if the expression of genes with IREs in *A. granulata* differs between iron-rich and iron-poor tissues. In the absence of iron, IRPs bind IREs. When this occurs in the 5’ UTR of an mRNA, ribosomes are blocked from translating the protein; thus, mRNAs with 5’ IREs will not be translated in the absence of free iron. When IRPs bind IREs in the 3’ UTR of an mRNA, they block endonucleases from degrading the mRNA, allowing multiple translations from a single mRNA molecule; thus, the translation of mRNAs with 3’ IREs is higher in the absence of free iron (Supplementary Figure 7). We compared the expression of genes with IREs between transcriptomes sequenced from the foot, girdle, ctenidia, and four developmentally-distinct regions of the radula from the same specimen of *A. granulata* we used for genome sequencing, and found expression of genes with IREs in all tissue types (Figure 3B). We quantified the expression of genes that contained 3’ IREs and 5’ IREs separately and found many genes with 5’ IREs had higher expression in the three anterior, iron-rich regions of the radula than the rest of the body (Supplemental Figure 8). Genes with 5’ IREs that are expressed in iron-rich tissues are likely transcribed into proteins in only those tissues because elsewhere in the body IRPs will bind to IREs and block translation. Consequently, the proteins encoded by these genes may be unique to the radula.

After identifying genes in *A. granulata* that contain 5’ IREs and are expressed at relatively high levels in the iron-rich anterior regions of the radula, we asked if these genes might have roles in the biomineralization of the radula. We used GO analysis to compare the molecular functions of the protein sequences coded by the genes with 5’ IREs to the protein sequences coded by the full set of genes from the *A. granulata* genome. We found that genes with a 5’ IRE that are highly expressed in the anterior of the radula are more likely to be associated with the molecular functions ‘response to inorganic substance’, ‘response to calcium ion’, and ‘response to metal ion’ (Supplementary Figure 9). This suggests that genes with a 5’ IRE that are highly expressed in the radula may be involved in the biomineralization of the apatite (calcium) cores of teeth and their magnetite (iron) caps. A previous study by Nemoto et al. identified a novel biomineralization protein (RTMP1) in the radula of another chiton (*Cryptochiton stelleri*), and proposed that RTMP1 played a role in iron biomineralization (Nemoto et al. 2019). We examined the mRNA of RTMP1 in *C. stelleri* and did not detect an IRE in either its 5’ or 3’ UTR. Thus there are genes that may be important to biomineralization in the chiton radula whose expression levels are not influenced by IREs.

### Two isoforms of ferritin may provide chitons with tissue-specific protection from oxidative stress

All metazoans require iron. However, free iron poses a threat to animals because it catalyzes the production of reactive oxygen species, which inflict damage on DNA and tissues (Dixon & Stockwell 2014). To transport iron safely, metazoans use the iron-binding protein ferritin. Previous work suggests that chitons use ferritin to transport iron to their radula (Kim et al. 1988). An iron response element (IRE) is present in the 5’ UTR of the heavy chain (or soma-like) ferritin that is expressed by all metazoans (Piccinelli & Samuelsson 2007-7). We found two isoforms of heavy chain ferritin in our gene models for *A. granulata:* a first isoform (isoform 1) that contains the conserved 5’ IRE, and a second isoform (isoform 2) that does not (Figure 4A).

**Figure 4:**
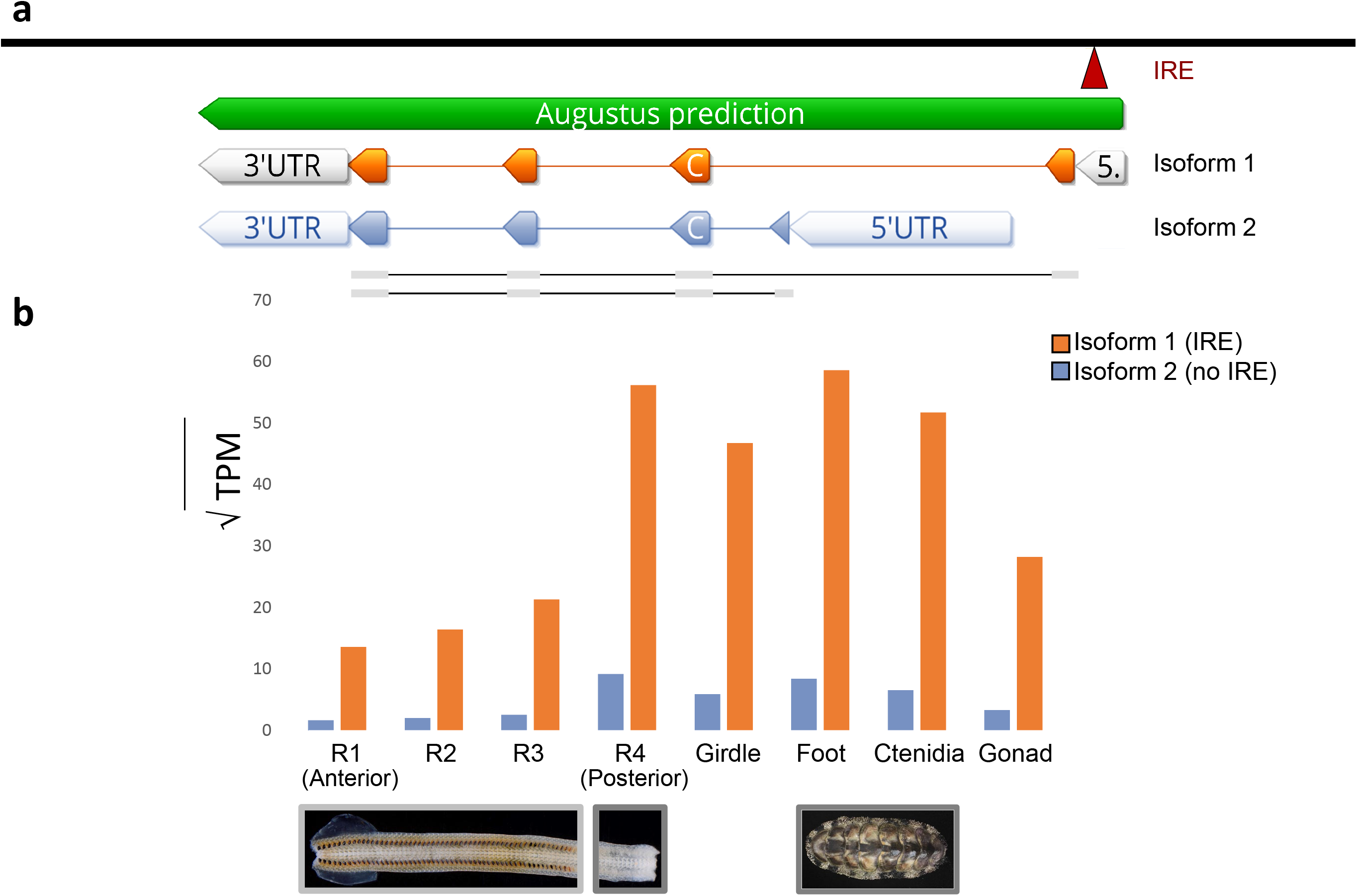
The two isoforms of heavy-chain ferritin recovered in A. granulata. (a) The locations of the transcription initiation sites and exons of isoform 1 of ferritin (orange, above) and isoform 2 of ferritin (blue, below). A 5’ IRE (red) is present in the 5’ untranslated region of isoform 1, but not in isoform 2. (b) Relative expression of both isoforms of ferritin across A. granulata tissues. The radula is divided into four developmentally distinct regions as in Figure 3. Isoform 1 is transcribed more highly throughout the body than isoform 2. Isoform 2 is transcribed at lower levels in the anterior (iron-rich) regions of the radula than in other tissues.

Isoform 1 of ferritin from *A. granulata* contains an IRE in the 5’ UTR, allowing this isoform to be translated only in the presence of free iron. By regulating the translation of ferritin, cells can transcribe ferritin mRNA continuously so they are primed to produce large quantities of ferritin protein rapidly if conditions require it. If no free iron is present, IRP will bind to the IRE and block translation. We found isoform 1 of ferritin is expressed at similar levels in all the transcriptomes we sequenced for *A. granulata*, including those for the foot, girdle, gonad, ctenidia, and all four regions of the radula (Figure 4B). Thus, when *A. granulata* needs to bind excess iron, it may be able to rapidly produce isoform 1 of ferritin protein throughout its body. We examined other mollusc genomes and transcriptomes and found a ferritin isoform present in all of them that is similar to *A. granulata* isoform 1 and contains the 5’ IRE.

Isoform 2 of ferritin in *A. granulata* lacks the 5’ IRE present in isoform 1. We identified an alternative transcription initiation site downstream of ferritin exon 1 in the *A. granulata* genome. Isoform 2 of ferritin, initiated at this downstream site, contains a different exon 1 than isoform 1 of ferritin, but shares exons 2-4 with isoform 1 (Figure 4). We examined other molluscan genomes and transcriptomes and did not find evidence for expression of a ferritin isoform similar to *A. granulata* isoform 2 (data available on Dryad). In *A. granulata*, transcripts of isoform 2 are expressed at a lower level than isoform 1 throughout all body tissues (foot, girdle, gonad, ctenidia) and in the posterior region of the radula that lacks iron mineralization (Figure 4B). Expression of isoform 2 is almost undetectable in the iron-rich regions of the radula. Without the 5’ IRE, translation of the mRNA of isoform 2 is not blocked in the absence of free iron. The 5’ IRE in ferritin is an important regulatory mechanism for protein production. In rats, for example, the expression of ferritin mRNAs is relatively constant across tissues but protein levels vary (Rogers & Munro 1987). Further, mutations in the 5’ IRE of ferritin cause hyperferritinaemia in mammals, an iron-related medical condition caused by an overproduction of ferritin protein (Thomson et al. 1999). We hypothesize that chitons use isoform 2 of ferritin to produce a low level of ferritin protein constitutively in tissues outside their radula as protection from the high concentrations of iron circulating through their bodies.

## Conclusions

The *A. granulata* genome is the first available genome for any chiton or any aculiferan. The information it provides improves our understanding of the evolution of biomineralization across Mollusca as well as lineage-specific innovations within chitons. Chitons are a valuable system for investigating biomineralization because they produce shell plates, sclerites, and iron-clad teeth. The unique combination of structures produced by chitons makes the *A. granulata* genome a resource for future studies of biomineralization. Although many genes involved in molluscan shell secretion are rapidly evolving (Jackson et al. 2006; Kocot et al. 2016), we were able to identify homologs of many conchiferan biomineralization genes in the *A. granulata* genome. The expression of several genes associated with conchiferan shell secretion in the girdle of *A. granulata* suggests these genes may function in sclerite biomineralization in chitons. This suggests a common underlying biomineralization mechanism for conchiferan shells and aculiferan sclerites, structures known to share some developmental pathways even though they arise via different cell lineages (Wollesen et al. 2017).

All metazoans require iron, but they must balance iron use against potential oxidative damage. Regulating iron is a particular concern for chitons because they biomineralize their teeth with magnetite. The genome of *A. granulata* contains more genes with iron response elements (IREs) than the genome of any other mollusc examined to date, indicating it has a larger proportion of genes regulated directly by iron. We identified two isoforms of ferritin in *A. granulata*, one that is iron-responsive and a second that is constitutively translated. We propose the second isoform of ferritin protects tissues outside the radula from oxidative stress by binding free iron. Chitons are an emerging model for studies of both biomineralization and iron homeostasis. The *A. granulata* genome will aid future studies by suggesting specific proteins and pathways to target with comparative studies of gene expression and gene manipulation.

## METHODS

### Specimen collection and preservation

We collected a single male specimen of *Acanthopleura granulata* from Harry Harris State Park in the Florida Keys (Special Activity License #SAL-17-1983-SR). We cut the majority of the foot into ~1 mm^2^ cubes and froze them at −80°C. We froze additional pieces of foot, girdle (dissected such that the tissue sample would not contain shell secretory tissue), ctenidia, gonad, and radula in RNAlater and stored them at −80°C as well.

### Genome and transcriptome sequencing

We extracted high molecular weight DNA from frozen samples of foot tissue from *A. granulata* using a CTAB-phenol chloroform method. We cleaned DNA for short read generation with the Zymo Clean and Concentrator Kit. For library preparation and sequencing, we sent cleaned DNA to the Genomics Services Lab at HudsonAlpha (Huntsville, AL), where it was sheared with a Covaris M220 to an average fragment size of 350 bp. These fragments were used to prepare an Illumina TruSeq DNA PCR-Free library, which was sequenced using one lane of an Illumina HiSeq X (2 × 150 bp paired-end reads).

For long-read sequencing, we cleaned DNA and enriched it for higher-molecular weight fragments by performing two sequential purifications using 0.4X AmPureXP magnetic beads. We generated long reads with four flow cells on an Oxford Nanopore Technologies GridION. We prepared two sequencing libraries with ligation kit LSK-108 and sequenced them on FloMin106 (R9.4.1) flow cells. We prepared the other two sequencing libraries with the updated ligation kit LSK-109 and sequenced them on R9.4.1RevD flow cells. We generated 2.19Gb, 4.41Gb, 7.87 Gb, and 8.4 Gb respectively across the four flow cells, for a total of 22.87 Gb, or >20x coverage with long-reads. Reads were basecalled with Guppy 4.0. We trimmed long reads with PoreChop (Wick 2018), which was set to remove chimeras (approximately 0.0005% of reads) and all residual adapter sequences.

To generate transcriptomes, we used the Omega Bio-tek EZNA Mollusc RNA Kit to extract RNA from girdle, ctenidia, gonad, foot, and four regions of radula (representative of visibly different stages of iron mineralization) of the same individual of *A. granulata* we used for genome sequencing. We synthesized and amplified complementary DNA (cDNA) from each tissue using the SmartSeq v4 Ultra Low-input RNA kit (Clontech) from 1 ng of input RNA with 17 cycles of PCR. We created eight dual-indexed sequencing libraries with the Illumina Nextera XT kit, using 1 ng of input cDNA. We sent the eight libraries to Macrogen (Seoul, South Korea) where they were pooled and sequenced on one lane of an Illumina HiSeq 4000 (2 x 100 bp paired-end reads).

### Genome and transcriptome assembly and quality assessment

We initially assembled the chiton genome with MaSuRCA v. 3.3.5 (Zimin et al. 2013), which consolidates paired-end data into super reads and then uses long-read data to scaffold and gap-fill. This produced an assembly with 2,858 contigs. We filtered and collapsed heterozygous contigs with Redundans v. 0.14a (Pryszcz & Gabaldón 07 08, 2016), decreasing the assembly to 1,285 contigs. To ensure that no contigs were incorrectly removed, we verified that all pre-Redundans contigs mapped to the post-Redundans assembly with bowtie2 (Langmead & Salzberg 2012); all contigs mapped and thus non-redundant data were not deleted. To help decontaminate reads and contigs, we used the Blobtools2 Interface to create blob plots (Blaxter & Challis 2018). Because Blobtools uses the NCBI nucleotide database to determine the identity of each scaffold, and chordate sequences vastly outnumber molluscan sequences in NCBI, Blobtools identified a large proportion of scaffolds as chordate. We identified contaminants as sequences that differed from the majority of scaffolds in both GC content and coverage and used BLAST to verify these sequences as bacterial before removing them from the assembly.

We scaffolded this reduced assembly with one lane of Bionano SAPHYR optical mapping, using two enzymes (BssSI and DLE1) and Bionano Solve v3.4’s scaffolding software, which resulted in 87 scaffolds. We ran REAPR v. 1.0.18 (Hunt et al. 2013), which map short read data and collect mapping statistics simultaneously, to determine accuracy of the assembly overall relative to all short-read data generated, and found despite reducing heterozygosity in the final assembly, 85.31% of paired-end reads map perfectly back to the genome assembly, indicating a complete genome assembly relative to the paired-end data.

To assess our genome assembly, we ran QUAST v. 5.0.2 (Gurevich et al. 2013). We assessed genome completeness with BUSCO v. 4.0.2 (Simão et al. 2015), using the proportions of nuclear protein-coding genes thought to be single-copy in the genomes of diverse metazoans (Metazoa odb9 dataset) and estimating the proportion of those that were complete, duplicated, fragmented, and absent.

We assembled the eight *A. granulata* transcriptomes with Trinity v. 2.84 (Grabherr et al. 2011), using the --trimmomatic and --normalize reads flags. We ran CD-Hit v. 4.8.1 (Fu et al. 2012) on each transcriptome separately to cluster isoforms. We also generated a composite transcriptome of all eight tissues (eight total transcriptomes including four separate radula regions) by combining reads and then following the same process described above. We used this composite transcriptome for annotation.

### Genome annotation

To annotate the *A. granulata* genome, we first generated a custom repeat library with RepeatModeler v. 2.0 (Smit & Hubley 2008-2015), which was used in all subsequent analyses. We trained MAKER v. 2.31.10 (Cantarel et al. 2008-1) on the composite transcriptome described above as well as predicted protein sequences from several other species of chitons that were generated previously (see Supplementary File 1). Using the highest quality gene models from the first as a maker-input gff3 (AED <0.5), we ran a second round of MAKER. From these resulting gene models, we used those with an AED <0.25 to train Augustus v3.0.3 (Stanke et al. 2006): we extracted gene models from the genomic scaffolds along with 1,000 bp of flanking sequence on either side to ensure complete genes, and ran them through BUSCO to produce an Augustus model (.hmm) file. Separately, we ran PASA 2.4.1 (Haas et al. 2003) on our composite transcriptome to maximize mapping transcripts to the genome assembly. We combined results from PASA and a trained Augustus run using the intersect tool in BEDtools v. 2.29.2 (Quinlan & Hall 2010), which removed identical sequences. This yielded a set of 81,691 gene models. When we ran a BUSCO v. 3.9 analysis (Metazoa odb9 dataset), we found a 15.2% duplication rate. To decrease duplications caused by transcripts predicted for the same locus by both Augustus and PASA that varied in length (and thus were not removed by the BEDtools intersect tool), we clustered the first set of gene models using cdhit-EST v. 4.8.1 (Fu et al. 2012), which we ran with the slow-but-accurate (-g) flag and with a cluster threshold value of 0.8. This produced a set of 20,470 genes. All commands we used are available in Appendix 1.

To identify annotated proteins in *A. granulata*, we first used Transdecoder (Douglas 2018) to produce peptide files of predicted proteins. We ran Interproscan on the set of 20,470 genes referenced above to identify GO terms and Pfam matches for proteins where possible. We used GHOSTX in the Kaas pipeline (Moriya et al. 2007) to identify KEGG pathways via comparisons to all the available molluscan taxa (*Lottia gigantea, Pomacea canaliculata, Crassostrea gigas, Mizuhopecten yessoensis*, and *Octopus bimaculoides*). Finally, we looked for shared GO terms between specific taxa with OrthoVenn(Xu et al. 2019), comparing *A. granulata* to *Lottia gigantea, Chrysomallon squaminiferon, Octopus bimaculoides*, and *Crassostrea gigas* (Supplementary Figure 10).

### Hox gene annotation and genomic comparisons

We located the Hox cluster of *A. granulata* by first creating a BLAST database of the *A. granulata* scaffolds and then querying this database with available chiton Hox sequences (Wanninger & Wollesen 2019). We marked *A. granulata* sequences with a BLAST hit at e-value 1e-8 as potential Hox sequences. We found one clear match for each previously identified chiton Hox gene, all in a single cluster within one scaffold. To verify the absence of *Post1*, we queried the *A. granulata* database with *Post1* sequences from five other molluscs (Wanninger & Wollesen 2019). All matched with low support to the existing *A. granulata Post2* sequence, so we concluded that *Post1* is absent from the *A. granulata* genome assembly.

To graphically examine synteny between *A. granulata* and other molluscan genome assemblies, we loaded each assembly and annotation into the online COGE SynMap2 (Haug-Baltzell et al. 2017) server and compared *A. granulata* to eight other annotated genomes with default SynMap2 settings. We exported dotplots for each pair of genomes to visualize syntenic regions (or lack thereof). Scaffolds in each dotplot were sorted by length, but differing assembly qualities made some dotplots difficult to read due to a high number of very small scaffolds. We assessed heterozygosity of several molluscan genomes and *A. granulata* by downloading raw paired-end data when possible and using GenomeScope2 online (Vurture et al. 2017).

To permit direct comparisons of repeat content within *A. granulata* and other molluscs, we ran RepeatModeler (Smit & Hubley 2008-2015) on the scaffolds of a subset of genome assemblies and *A. granulata*. We used the same default parameters for each run and quantified the number of elements in each repeat family identified by RepeatModeler for each genome assembly we analyzed (LINEs, SINEs, etc.; Supplementary Table 7).

### Orthology inference

To identify orthologous genes shared between *A. granulata* and other molluscs, we used OrthoFinder v. 2.3.7 (Emms & Kelly 2015). We analyzed three separate sets of data: 1) *A. granulata* and genomes of nineteen other lophotrochozoans, including fourteen other molluscs, two annelids, one brachiopod, one phoronid, and one nemertean 2) *A. granulata* and a subset of molluscan genomes for detailed comparisons of biomineralization genes and; 3) *A. granulata* and an expanded set of data including both genomes and transcriptomes, including several transcriptomes from aculiferans other than *A. granulata.* For all three analyses we used the unclustered 81,691 gene set for *A. granulata*, knowing that duplicated gene models would cluster together. We removed sequences from our orthogroups that were identical to longer sequences where they overlapped, as well as fragmented sequences shorter than 100 amino acids, using uniqHaplo(Anon). We retained orthogroups that had a minimum of four taxa, aligned the sequences within them with MAFFT (Katoh et al. 2002), and cleaned mistranslated regions with HmmCleaner (Di Franco et al. 2019). We used AlignmentCompare (https://github.com/kmkocot/basal_metazoan_phylogenomics_scripts_01-2015) to delete sequences that did not overlap with all other sequences by at least 20 AAs (starting with the shortest sequence meeting this criterion).

### Phylogenetic analyses

For species tree reconstruction, in cases where two or more sequences were present for any taxon in a single-gene alignment, we used PhyloPyPruner 0.9.5 (https://pypi.org/project/phylopypruner/) to reduce the alignment to a set of strict orthologs. This tool uses single-gene trees to screen putative orthogroups for paralogy. To build single-gene trees based on orthologs, we trimmed alignments with BMGE v1.12.2 (Criscuolo & Gribaldo 2010) and constructed approximately maximum likelihood trees for each alignment with FastTree2(Price et al. 2010) using the “slow” and “gamma” options. We then used these alignments in PhyloPyPruner with the following settings: --min-len 100 --min-support 0.75 --mask pdist --trim-lb 3 --trim-divergent 0.75 --min-pdist 0.01 --trim-freq-paralogs 3 --prune MI. For datasets 1 (“genomes”) and 3 (“all_taxa”), only orthogroups sampled for at least 85% of the total number of taxa were retained for concatenation. For dataset 2 (“biomin_subset”), only orthogroups sampled for all eight taxa were retained. Phylogenetic analyses were conducted on the supermatrix produced by PhyloPyPruner v. 1.0 in IQ-TREE v. 1.6.12 (Nguyen et al. 2015) using the PMSF model (Wang et al. 2018) with a guide tree based on the LG model. Topological support was assessed with 1,000 rapid bootstraps.

### Screening for known biomineralization genes

We identified known molluscan biomineralization genes of interest in the chiton genome by first making a BLAST protein database of protein sequences for the 81,691-gene model set of *A. granulata* annotations, translated by Transdecoder (Grabherr et al. 2011; Douglas 2018). We then used the highest-quality protein sequence of that gene available on NCBI (complete where available, longest if only incomplete protein sequences existed) as a query for each biomineralization gene of interest with an initial e-value cutoff of 1e-8. In cases where multiple hits of similar support resulted, we selected the correct hit by constructing a phylogeny in RAxML v. 8.2.12 (Stamatakis 2014) under the GTRGAMMA model with rapid bootstrapping and a best-scoring maximum-likelihood tree search in one run, with the number of bootstrap replicates determined by majority-rule consensus (autoMRE). This produced a list of sequences from *A. granulata* that matched the biomineralization genes of interest, and allowed us to narrow down our list of potential biomineralization genes present in *A. granulata*.

We used the above set of gene queries from other molluscs to identify the ortholog group from the above OrthoFinder2 on the subset of genomes selected as biomineralization representatives across Mollusca. We used the complete CDS or longest mRNA for each gene as a nucleotide query to search our orthogroups, again with an e-value cutoff of 1e-8 to identify the orthogroup(s) likely contained that particular biomineralization gene of interest. This produced a list of orthogroups that contained sequences with high similarity to the query, often multiple orthogroups per gene (Supplementary Table 4). This was expected due to clustering within OrthoFinder2. We used NCBI BLAST to verify the identity of the orthologous gene sequences by verifying that the top hits for each in BLAST matched the biomineralization gene of interest. We then examined these orthogroups to locate the previously identified *A. granulata* gene model that matched to each biomineralization protein. The query sequences for each gene sought in *A. granulata* are available in Supplementary Table 8.

Silk-like proteins share similar amino acid composition throughout Metazoa, but the genes that code for them are difficult to identify in genomes because their highly-repetitive sequences are often missed by traditional gene annotation tools (McDougall et al. 2016). We looked for silk-like proteins with SilkSlider (McDougall et al. 2016), run with default settings but using SignalP v. 4.01 (Nielsen 2017), which identifies potential silk-like proteins by locating low-complexity repetitive domains and signal peptides. The 31 proteins identified as silk-like by SilkSlider were then uploaded to the SignalP 5.0 webserver (Almagro Armenteros et al. 2019) for further predictions of signal peptides associated with extracellular localization.

To locate and quantify iron response elements (IREs), we screened the 20,470-gene *A. granulata* gene model set using the SIREs 2.0 (Campillos et al. 2010) web server. We also ran SIRE on the subset of genomes used for biomineralization analyses (see OrthoFinder above) for comparison. We compensated for differences in annotation methods by first clustering all coding sequences from each genome with CD-Hit-EST (Fu et al. 2012) with a cluster threshold of 0.8 (to match the threshold value we used earlier to reduce redundancy in the annotations of the *A. granulata* genome). We then ran SIRE on each of these sets of predicted transcripts. We only accepted predicted IREs scored as “high quality” according to the SIRE metric (indicating both sequence and structural characteristics of a functional IRE). We pulled chiton genes containing a high quality IRE from the eight different tissue transcriptomes generated for genome annotation and assessed expression by mapping each back to the genome with Salmon v. 0.11.3 (Patro et al. 2017-4) to generate quantifications of reads per transcript, and running these quantifications through edgeR (Robinson et al. 2010) to account for transcript length (TPM) and permit direct comparisons of gene expression. We also separated 3’ and 5’ IREs by subsetting the high quality IREs based on whether the IRE was located at the beginning or end of the sequence. We made heatmaps with log-transformed data to compensate for outliers in expression levels with R package prettyheatmap (Kolde 2012). We then analyzed the GO terms enriched in the separate sets of 5’- and 3’-containing genes that were highly expressed in the radula with GOrilla (Eden et al. 2009), using the complete protein set as a background dataset and the sets of IRE-containing genes as the target list.

## Supplementary Material

The West Indian Fuzzy Chiton *Acanthopleura granulata* genome and transcriptomes from the same individuals have been deposited in the NCBI database as BioProject PRJNA578131. The genome project is registered in NCBI as JABBOT000000000. All raw reads for both the genome and transcriptomes are available online at the NCBI Sequence Read Archive under the same BioProject PRJNA578131. Transcriptome data of other species of chitons used for genome annotation are available online at the NCBI Sequence Read Archive under BioProjects PRJNA626693 and PRJNA629039. The genome assembly, all sets of gene models discussed in the manuscript, functional annotations, and supporting documentation for the biomineralization genes described are available in Dryad with the identifier doi:X [For the purposes of peer review, data are currently available via Box: https://alabama.box.com/s/1hsryfff61i01qrljyasrjnu8j7qg2nj. All code used in this study is available in Supplementary Material, Appendix 1.

## Acknowledgements

We thank Ken Halanych and the crew and scientists of the Icy Inverts cruises aboard R/V Lawrence M. Gould and R/V Nathaniel B. Palmer for collecting *Callochiton* sp., Christoph Held, the Alfred Wegener Institute, and the scientists and crew of the PS96 cruise aboard R/V Polarstern for facilitating collection of *Nuttallochiton* sp., Julia Sigwart and Lauren Sumner-Rooney for collecting *Leptochiton asellus*, and Alexandra Kingston and Daniel Chappell for collecting *Chiton marmoratus and Chiton tuberculatus*, all used for transcriptome sequencing. CM was supported by a Malacological Society of London grant for transcriptome sequencing of *Acanthopleura gemmata*. We thank Kerry Roper for RNA QC of the *Acanthopleura gemmata* transcriptome samples. We thank the staff of the Smithsonian Marine Station at Ft. Pierce and the Keys Marine Lab for providing housing during collections. RMV thanks John Sutton, Michael McKain, and Michelle Lewis for assistance in optimizing nanopore library preparation protocols.

