## Supplementary Figures/Tables/Document for "The iron-responsive genome of the chiton *Acanthopleura granulata*"

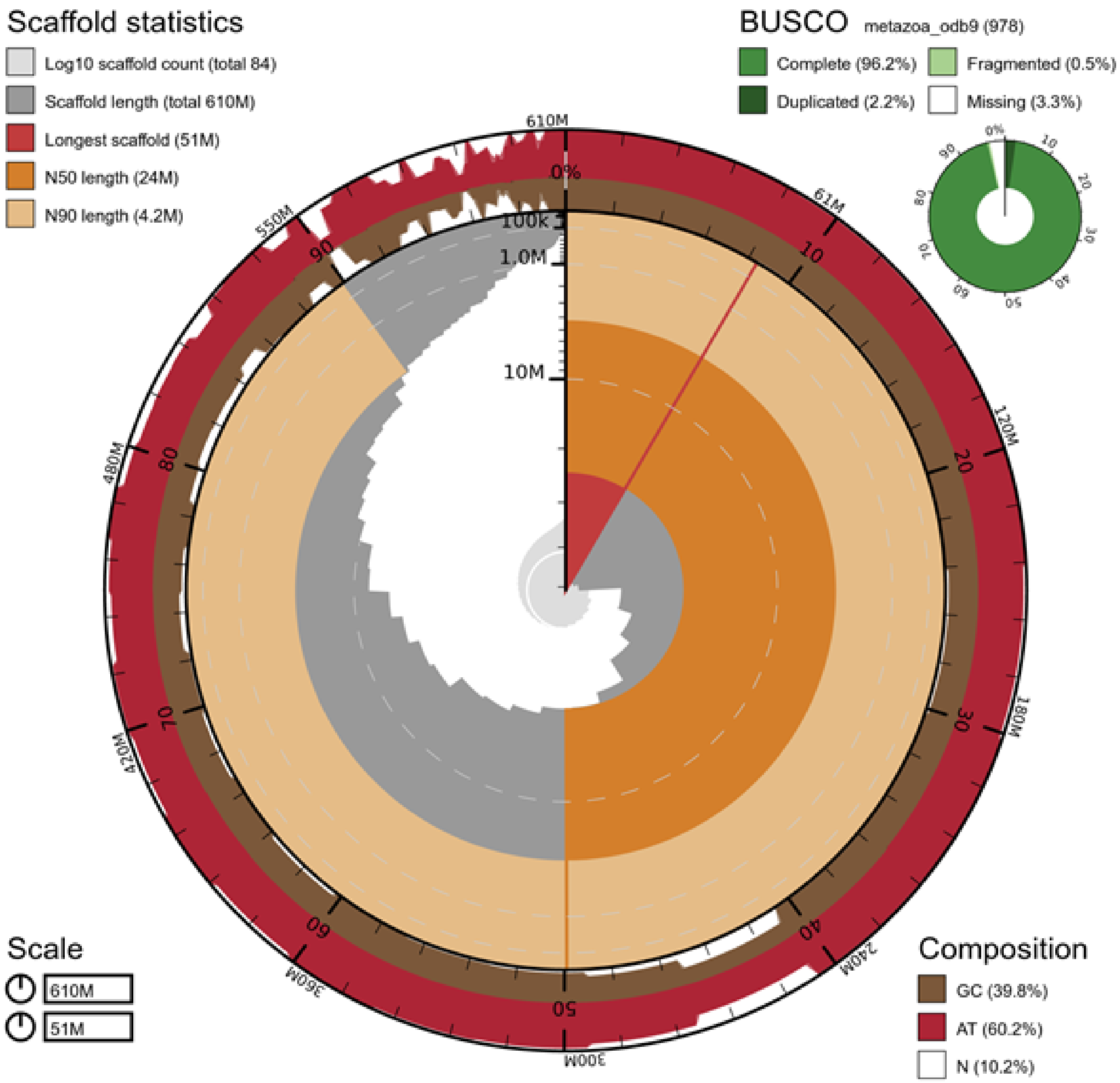


**Supplementary Figure 1:** Snail plot of the *Acanthopleura granulata* genome assembly. N50 is shown in orange. A preliminary BUSCO analysis (BUSCO v.3, gene set metazoan obd9) in the upper right indicates 96.2% completeness- with updates to BUSCO this improved to 97.4% completeness.

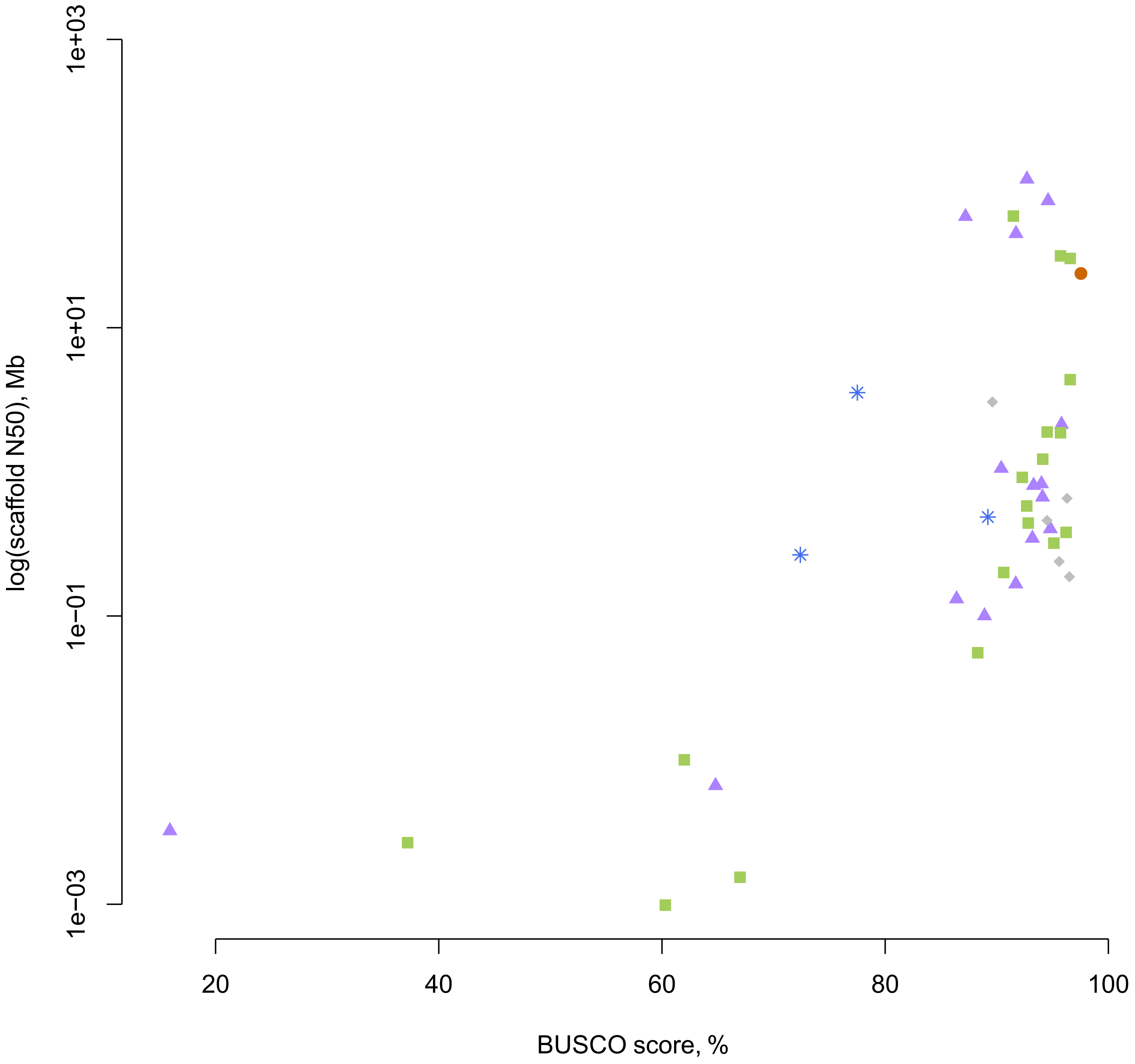


**Supplementary Figure 2:** An illustration of the quality of available molluscan genomes as a scatter plot of BUSCO completeness score and a log transform of the N50 of scaffolds in megabases for each genome. The *A. granulata* genome is the orange circle, bivalves are purple triangles, gastropods are green squares, and cephalopods are blue stars. Grey diamonds are representative genomes of other lophotrochozoans (specifically those used as outgroups in the homolog gene searches in this study).

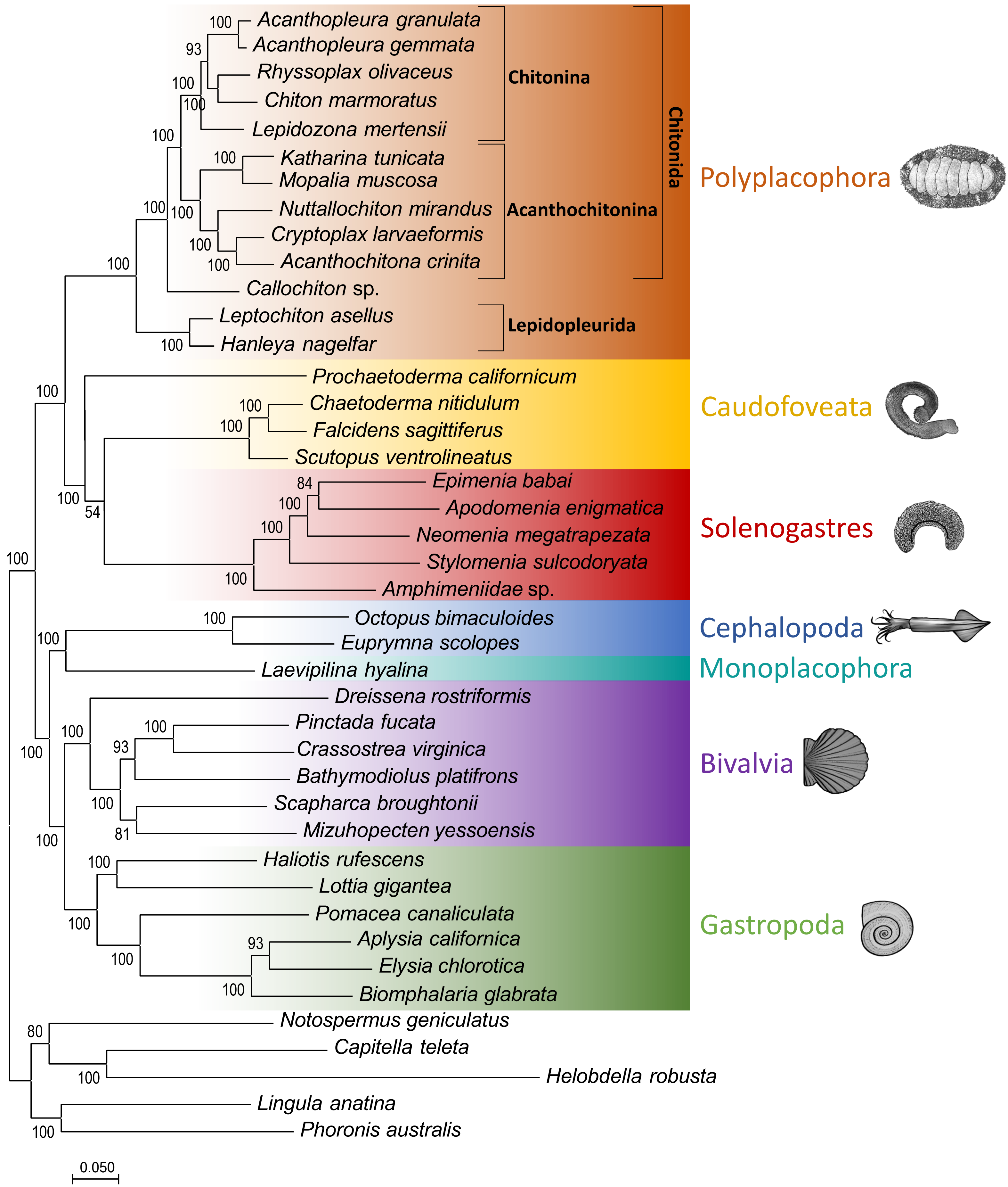


**Supplementary Figure 3:** An expanded phylogeny based on transcriptome as well as genome sequences, showing the position of A. granulata within a broader range of chitons and other molluscs. Aculifera (Aplacophora [Caudofoveata + Solenogastres] + Polyplacophora) was recovered with maximal support. Within Polyplacophora, A. granulata is recovered within Chitonina, which is also maximally supported.

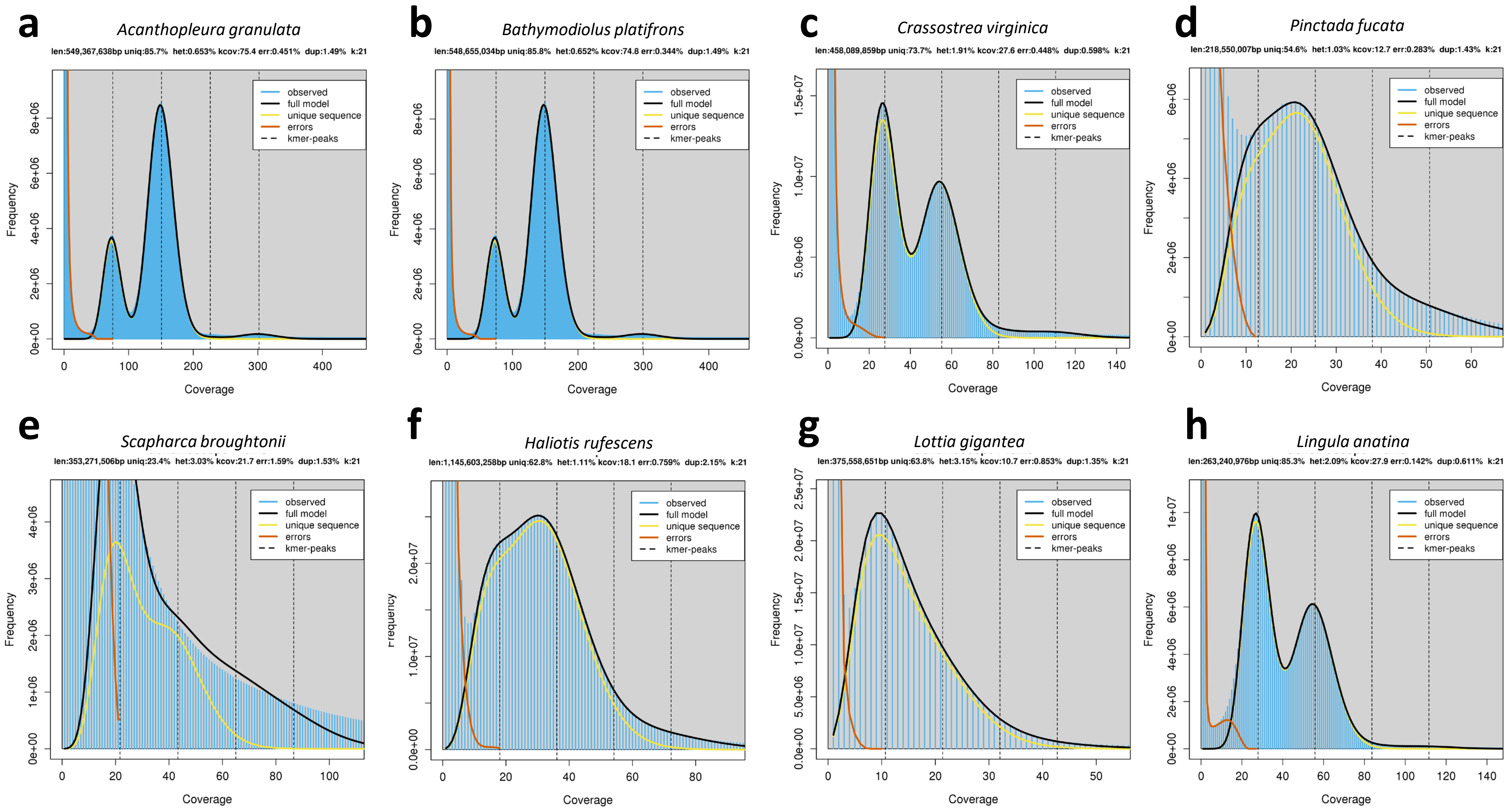


**Supplementary Figure 4:** Genome Scope analyses of the paired-end data for available molluscan genomes. Heterozygosity is measured via k-mer distribution and available at the top of each graph as het: %.

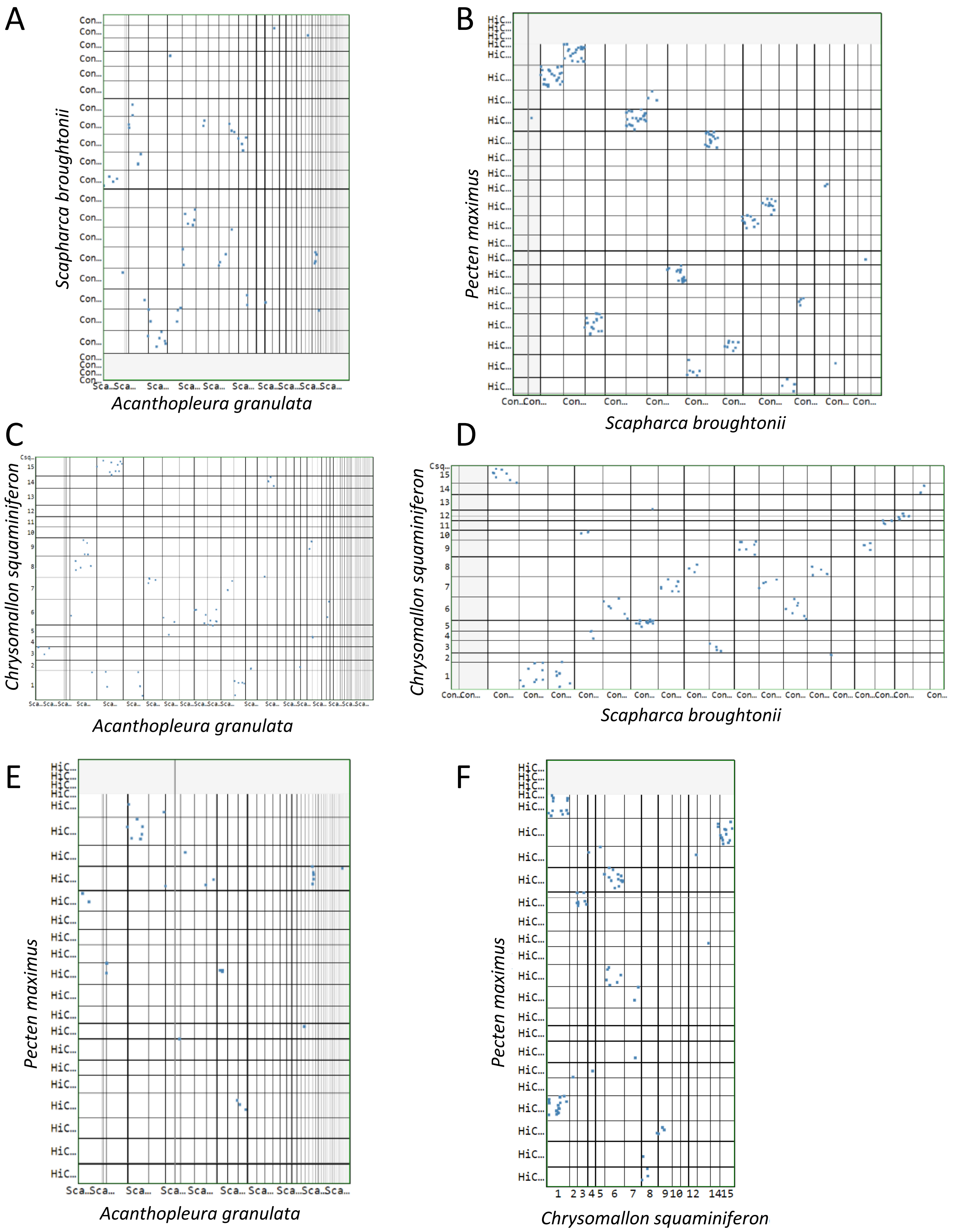
A
Scapharca broughtonii
Acanthopleura granulata
B
Pecten maximus
Scapharca broughtonii
C
Chrysomallon squaminiferon
Acanthopleura granulata
D
Chrysomallon squaminiferon
Scapharca broughtonii
E
Pecten maximus
Acanthopleura granulata
F
Pecten maximus
Chrysomallon squaminiferon

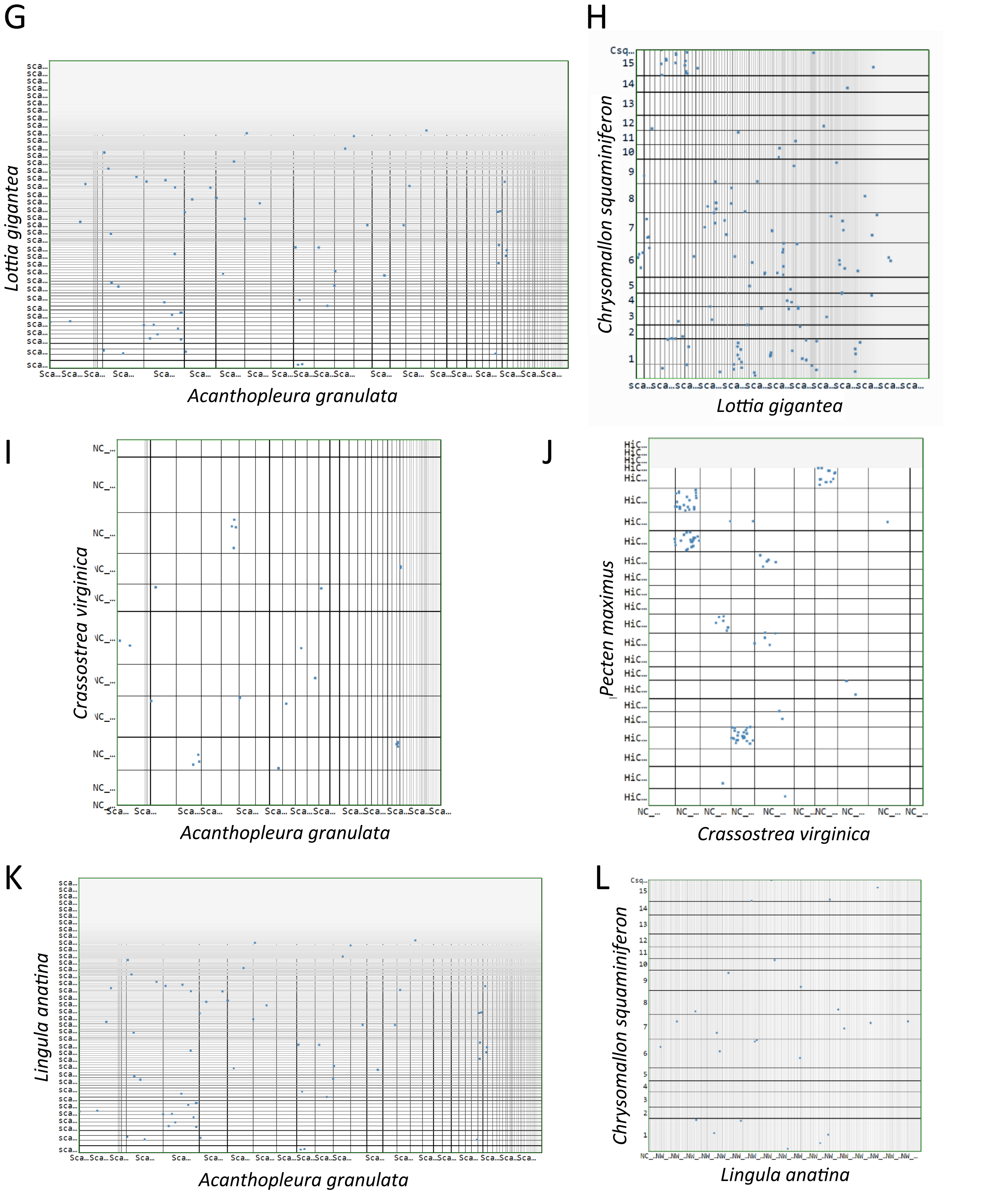


**Supplementary Figure 5:** Comparisons of synteny between the genome of *Acanthopleura granulata* and other lophotrochozoans, compared to synteny between conchiferans. Blue dots represent regions of synteny as generated in SynMap. Regardless of assembly quality, there is more synteny between two conchiferans (B,D,F,H,J) than between chonchiferans and *A. granulata* (A,C,E,G,I), and *A. granulata* does not share more synteny with a brachiopod than any conchiferan (K-L).

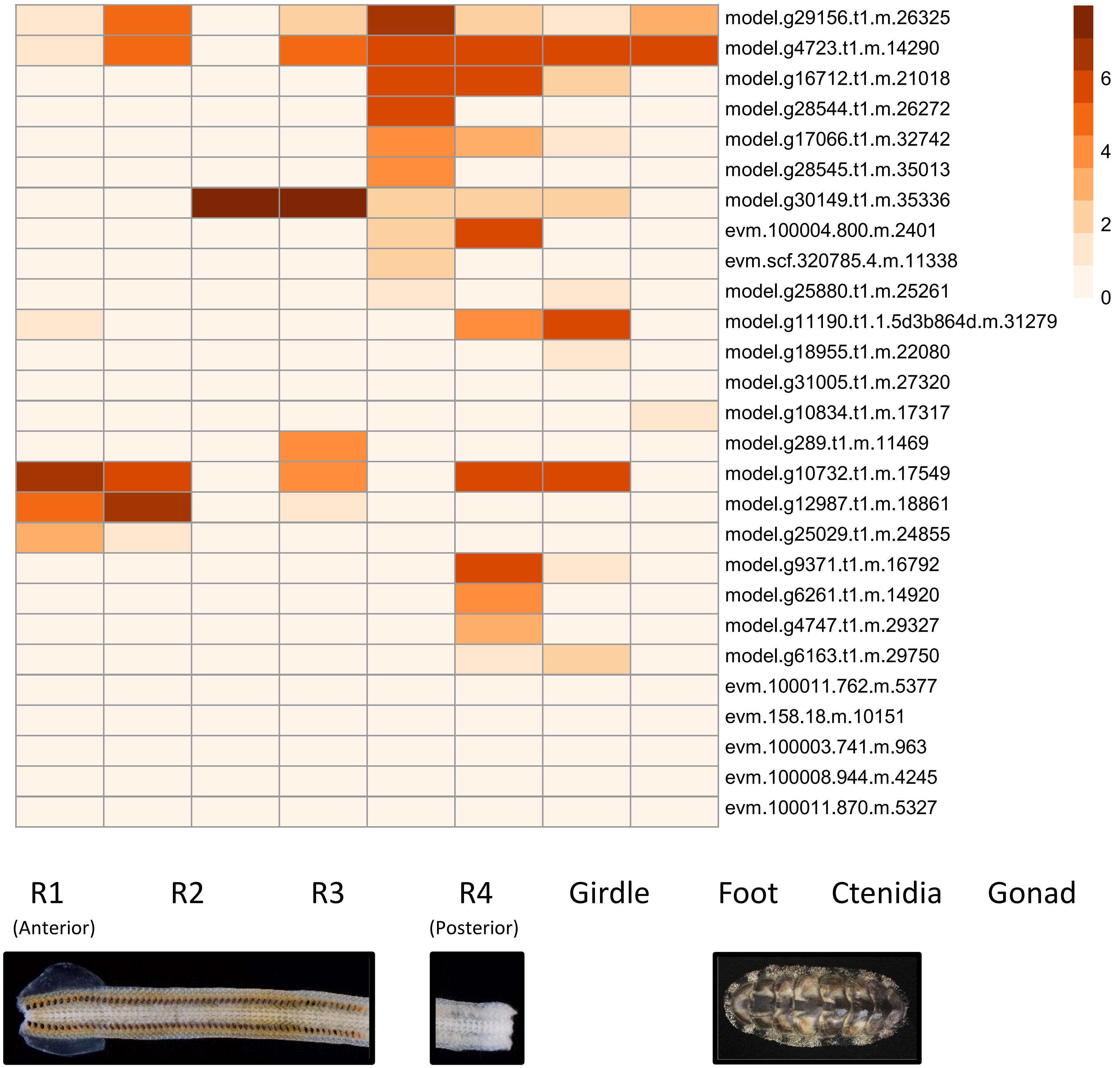


**Supplementary Figure 6:** Heat map of the expression (log(TPM)) of the 31 genes identified as potentially silk-like by SilkSlider. Expression is higher for several of these genes in the girdle and radula, supporting a potential role of these genes in biomineralization.

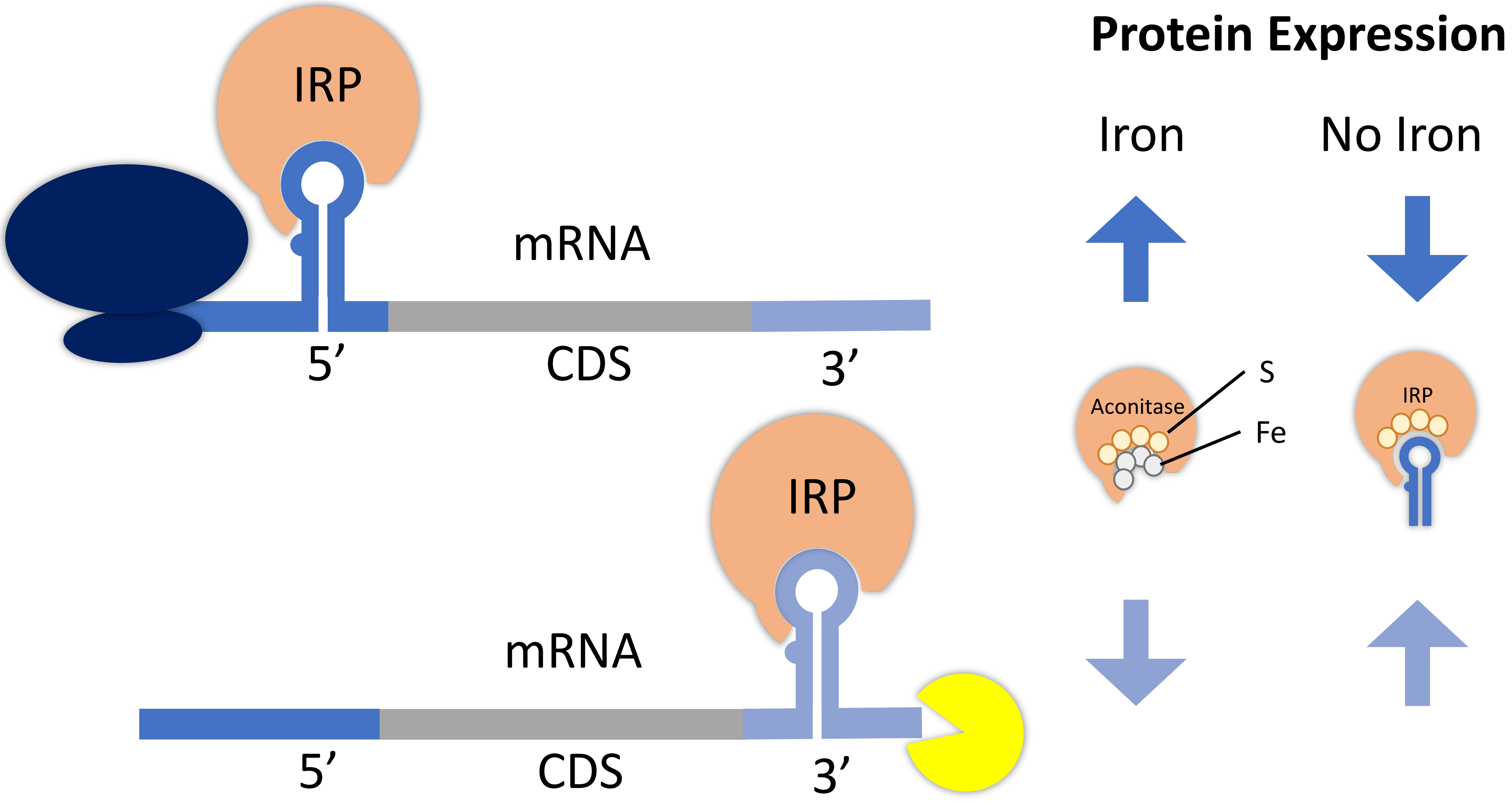


**Supplementary Figure 7:** The mechanism through which iron response elements (IREs) function. When free iron is present, an IRP will not bind to an IRE because iron (Fe) binds to sulfur (S) at the enzyme’s active site and changes the enzyme’s conformation such that it functions as aconitase (a TCA cycle participant). In the absence of free iron, IRPs bind to IREs, affecting translation rates. When IRPs bind to IREs in the 5’ UTR of an mRNA, they block ribosomes (navy) and prevent translation; thus, mRNAs with 5’ IREs will be translated in the presence of free iron. When IRPs bind to IREs in the 3’ UTR, they block endonucleases (yellow) from degrading mRNA, thereby allowing multiple translations from a single mRNA molecule; thus, the amount of protein produced from mRNAs with 3’ IREs will decrease in the presence of free iron.

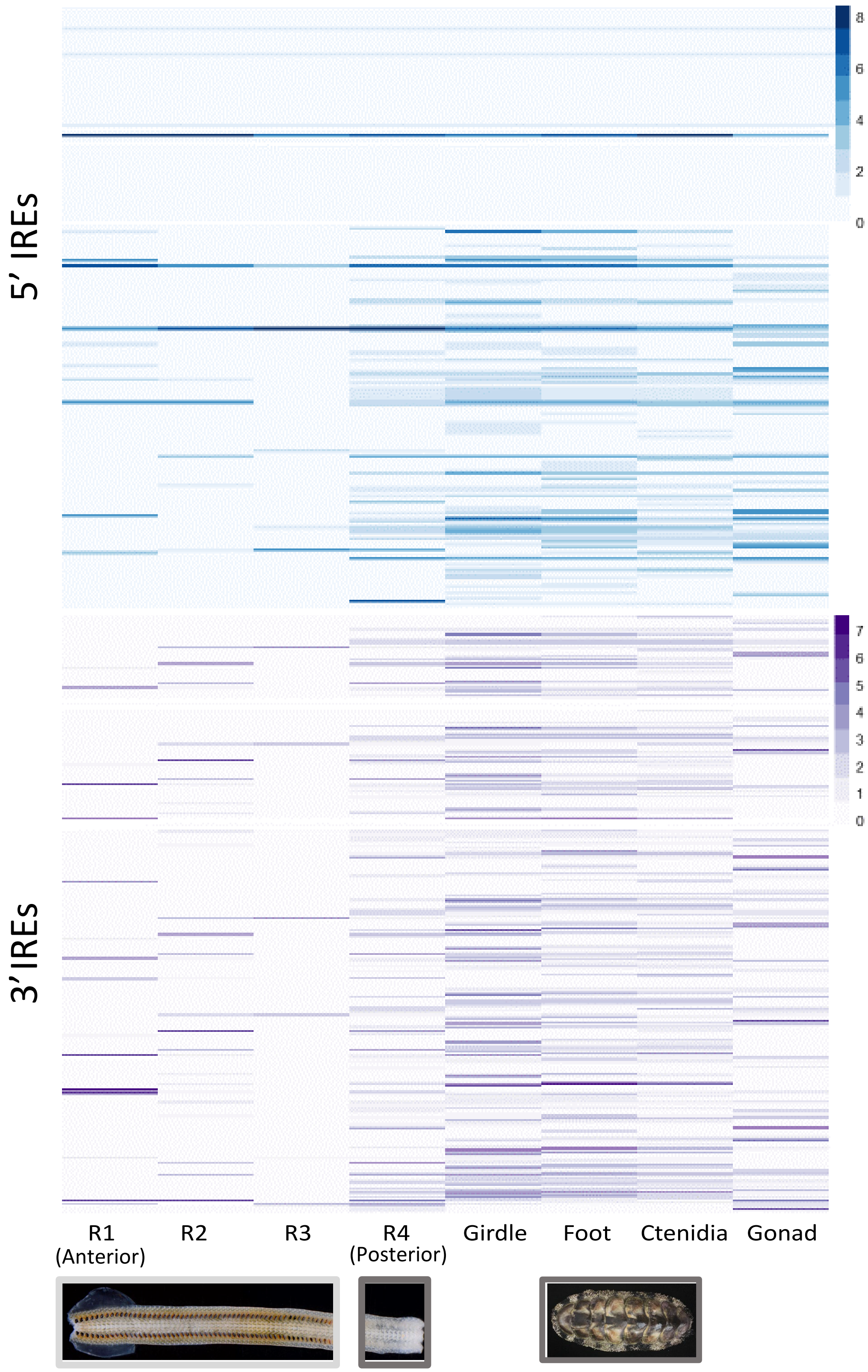


**Supplementary Figure X:** The relative expression [log10(TPM)] of transcripts containing 5' IREs (upper, blue) and 3' IREs in the different tissues of A. granulata. The radula is divided into four developmentally distinct regions: R1,the most anterior region, contains teeth used for feeding; R2 contains teeth that are developed but are not yet used for feeding; R3 contains developing teeth that contain iron oxide; and R4, the most posterior region, contains developing teeth that have yet to be coated with iron. We found a greater number of 5' IRE-containing genes than 3' IRE-containing genes that are more highly expressed in the anterior sections of the radula (R1-R3) than in the remaining body tissues.

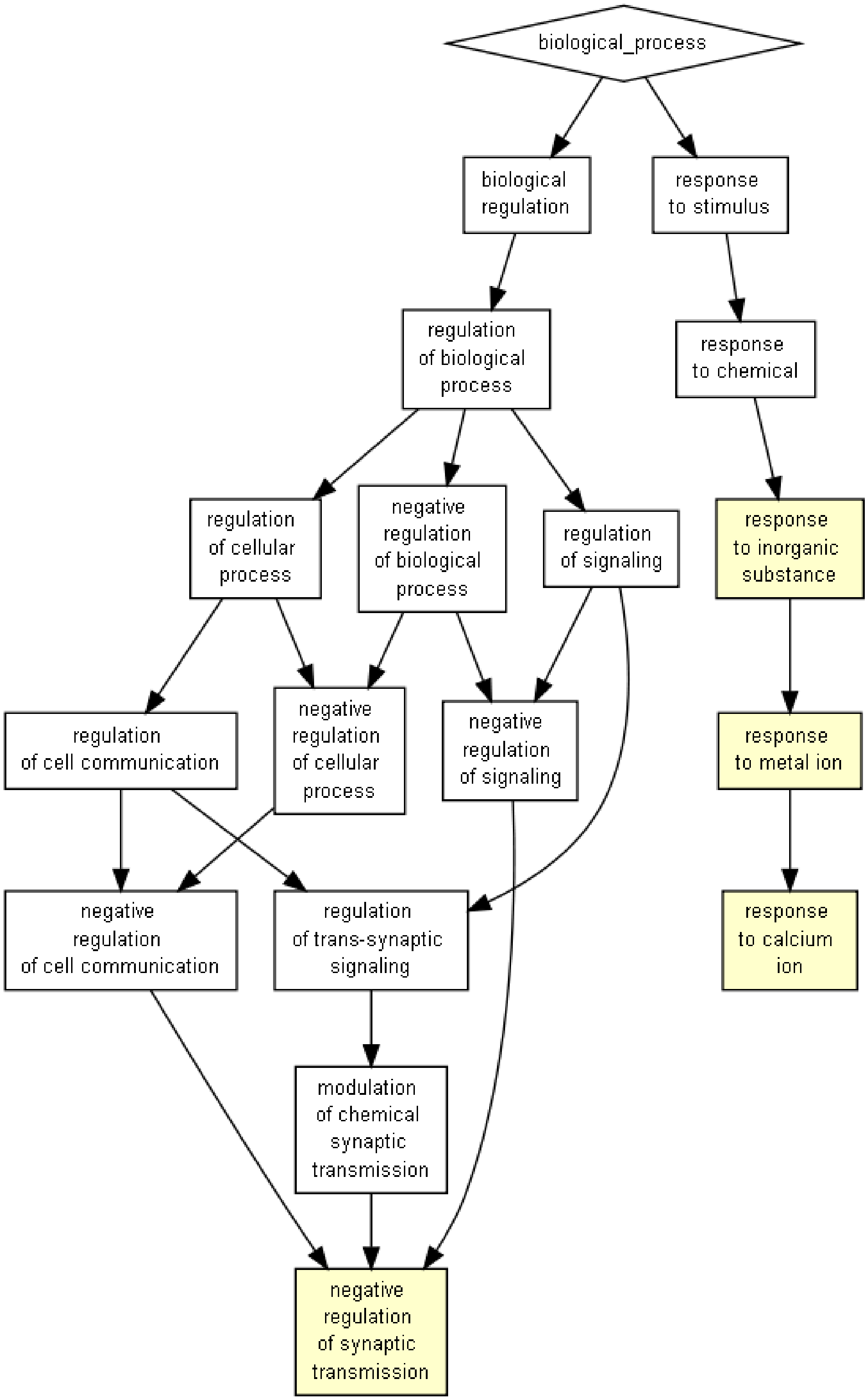


**Supplementary Figure 9:** GO-term enrichment in the upregulated genes of *Acanthopleura granulata* that contain 5' IREs. Yellow colored boxes indicate biological process GO terms that are enriched in these genes relative to a background gene set of all *A. granulata* genes. The genes that are highly expressed in the radula and have 5'-IREs are enriched for response to inorganic substance, response to metal ion, and response to calcium ion.

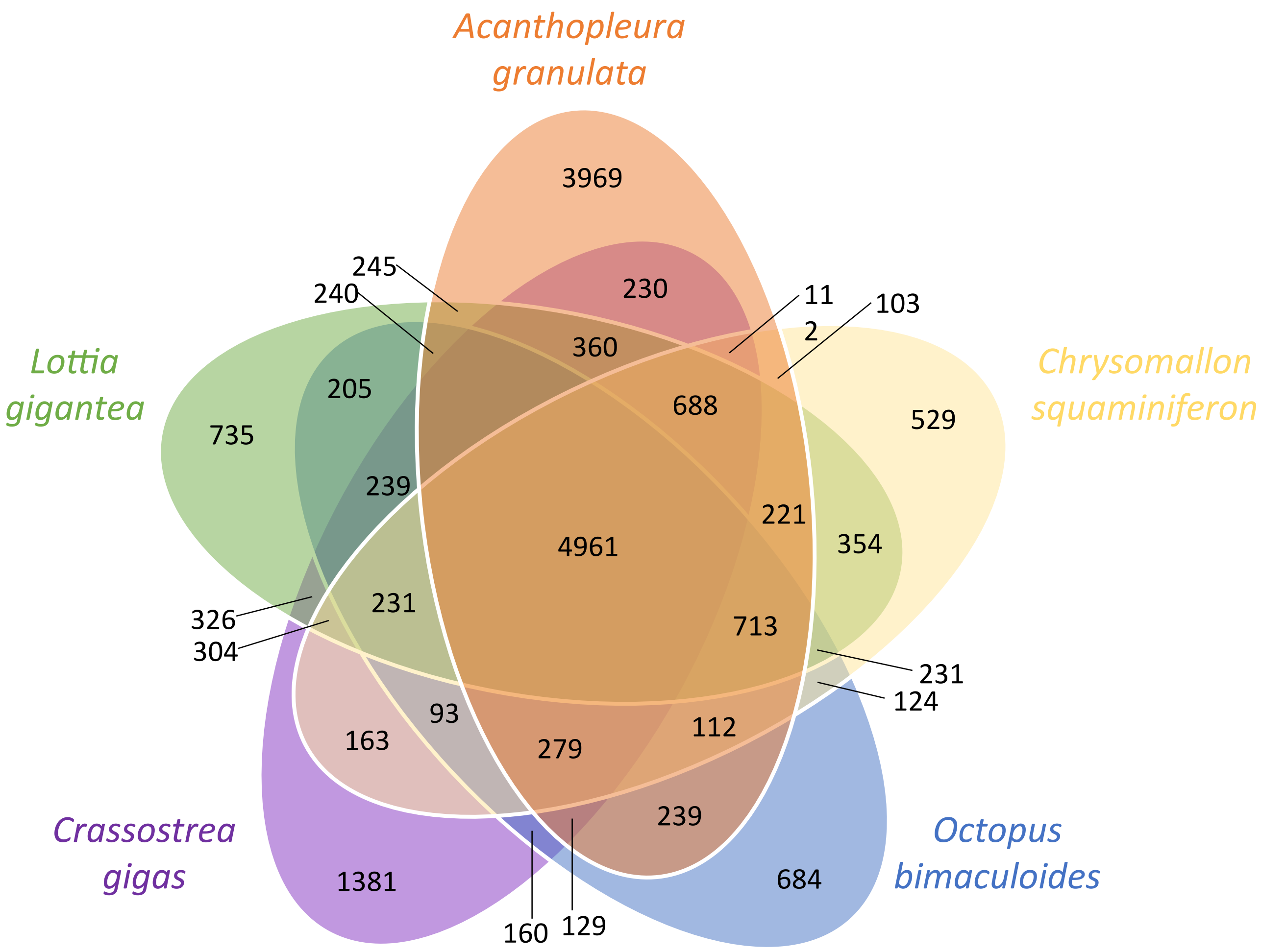


**Supplementary Figure 10:** Venn diagram of the overlap of orthologous proteins between five molluscs, including *Acanthopleura granulata. A. granulata* has almost as many unique proteins as it shares will all other molluscs present, highlighting the divergenge between *A. granulata* and other sequenced conchiferans.

**Supplementary Table 1:** Table of lophotrochozoan genomes available at the time of publication. Details of scaffold continuity were calculated from each genome with QUAST. Completeness scores (BUSCO) were calculated from each genome assembly with BUSCO 3.1, using metazoan gene set obd9. For genomes with available gene models, completeness scores were also calculated with BUSCO 3.1, metazoan gene set obd9, in -transcriptome mode.

| Clade | Taxon | Length (Mb) | Number of Scaffolds | N50 (Mb) | Longest scaffold (Mb) | BUSCOgeno(c) | BUSCOtrans(c) | Citation | Source |
|---|---|---|---|---|---|---|---|---|---|
| Polyplacophora | *Acanthopleura granulata* | 606 | 87 | 24.00 | 50.9 | 97.4 | 96.9 | Current study | NCBI PRJNA578131 |
| Cephalopoda | *Euprymna scolopes* | 5710 | 59,146 | 3.55 | 29.7 | 77.5 | 87.9 | Belcaid et al. 2019 | NCBI PRJNA470951 |
| Cephalopoda | *Octopus bimaculoides* | 2338 | 151,674 | 0.49 | 4.1 | 89.2 | 92.8 | Albertin et al. 2015 | NCBI PRJNA305125 |
| Cephalopoda | *Octopus vulgaris* | 1349 | 77,681 | 0.27 | 3 | 72.4 | none | Zarrella et al 2019 | NCBI PRJNA492973 |
| Bivalvia | *Argopecten pupuratus* | 708 | 89,727 | 1.06 | 11.13 | 90.4 | 90.9 | Li et al 2018 | http://gigadb.org/dataset/view/id/100419 |
| Bivalvia | *Bathymodiolus platifrons* | 1658 | 65,664 | 0.35 | 2.79 | 93.2 | 86.6 | Sun et al. 2017 | NCBI PRJNA328542 |
| Bivalvia | *Chlamys farreri* | 910 | 388,151 | 0.67 | 6.57 | 94.1 | none | Li et al 2017 | NCBI PRJNA185465 |
| Bivalvia | *Crassostrea gigas* | 558 | 7,658 | 0.40 | 1.96 | 94.8 | 83.4 | Zhang et al. 2012 | NCBI PRJNA276446 |
| Bivalvia | *Crassostrea virginica* | 685 | 11 | 75.94 | 104.16 | 94.6 | 98.4 | Gomez-Chiarri et al 2015 | NCBI PRJNA379157 |
| Bivalvia | *Dreissena polymorpha* | 1798 | 144 | 107.57 | 190.5 | 92.7 | none | Penarrubia er al 2015 | https://zebra_mussel.s3.msi.umn.edu/Dpolyporpha_Assembly.V2.Final_wMito.fasta.gz |
| Bivalvia | *Dreissena rostriformis* | 1241 | 18,504 | 0.13 | 1.15 | 86.4 | 81 | Calcino et al 2019 | https://phaidra.univie.ac.at/view/o:980132 |
| Bivalvia | *Mizuhopecten yessoensis* | 988 | 82,659 | 0.83 | 7.5 | 94 | 98.7 | Wang et al. 2017 | NCBI PRJNA390633 |
| Bivalvia | *Modiolus philippinarum* | 2630 | 74,575 | 0.10 | 0.715 | 88.9 | 73.5 | Sun et al. 2017 | NCBI PRJNA328544 |
| Bivalvia | *Mytilus galloprovincialis* | 1399 | 1,002,334 | 0.00 | 0.06 | 15.9 | 4.2 | Murgarella et al 2016 | NCBI PRJNA262617 |
| Bivalvia | *Pinctada fucata* | 815 | 29,306 | 0.17 | 1.26 | 91.7 | 79.4 | Takeuchi et al. 2016 | http://marinegenomics.oist.jp/pinctada_fucata |
| Bivalvia | *Pinctada imbricata* | 991 | 5,039 | 59.03 | 104.6 | 87.2 | none | Du et al 2017 | NCBI PRJNA283019 |
| Bivalvia | *Scapharca broughtonii* | 884 | 1026 | 44.99 | 55.6 | 91.72 | 90.6 | Bai et al. 2019 | NCBI PRJNA521075 |
| Bivalvia | *Venustaconcha ellipsiformis* | 1590 | 371,427 | 0.01 | 0.31 | 64.8 | none | Renault 2018 | NCBI PRJNA433387 |
| Bivalvia | *Saccostrea glomerata* | 784 | 10,107 | 0.80 | 7.15 | 93.3 | 91.5 | Powel et al 2018 | NCBI PRJNA414259; http://soft.bioinfo-minzhao.org/srog/# |
| Bivalvia | *Lutraria rhynchaena* | 544 | 622 | 2.14 | 13.02 | 95.8 | none | Thai et al 2019 | NCBI PRJNA548223 |
| Gastropoda | *Aplysia californica* | 927 | 4,332 | 0.92 | 0.61 | 92.3 | 98.6 | Broad Institute | NCBI PRJNA13635 |
| Gastropoda | *Biomphalaria glabrata* | 916 | 331,401 | 0.06 | 2.18 | 88.3 | 92.3 | Adema et al. 2017 | NCBI PRJNA290623 |
| Gastropoda | *Conus tribblei* | 2160 | 1,126,156 | 0.00 | 0.08 | 37.2 | none | Barghi et al 2016 | NCBI PRJNA285711 |
| Gastropoda | *Cumia reticulata* | 44 | 144,469 | 0.00 | 0.016 | 67 | none | Modica MV et al 2015 | NCBI PRJEB9058 |
| Gastropoda | *Elysia chlorotica* | 557 | 9,989 | 0.44 | 2.35 | 92.8 | 96.9 | Bhattacharya et al 2013 | NCBI PRJNA484060 |
| Gastropoda | *Haliotis discus hannai* | 1865 | 80,032 | 0.20 | 2.21 | 90.6 | 33.6 | Nam et al. 2017 | NCBI PRJNA317403 |
| Gastropoda | *Haliotis rubra* | 1378 | 2,854 | 1.23 | 11.18 | 94.1 | none | Kijas et al 2019 | NCBI PRJNA489521 |
| Gastropoda | *Haliotis rufescens* | 1498 | 8,371 | 1.90 | 13.19 | 94.5 | none | Masonbrink et al 2019 | https://abalone.dbgenome.org/downloads |
| Gastropoda | *Lottia gigantea* | 360 | 4,475 | 1.87 | 9.39 | 95.7 | 96.5 | Simakov et al. 2012 | NCBI PRJNA259762 |
| Gastropoda | *Lymnaea stagnalis* | 411 | 85,244 | 0.01 | 0.095 | 62 | none | Davison et al 2017 | NCBI PRJEB11470 |
| Gastropoda | *Patella vulgata* | 28 | 29,489 | 0.00 | 0.015 | 60.3 | none | Kenny et al 2015 | https://ora.ox.ac.uk/objects/uuid:6471e7d4-dd34-4eb3-883f-a5c438f11731 |
| Gastropoda | *Pomacea canaliculata* | 440 | 24 | 31.53 | 45.4 | 95.7 | 98.7 | Liu et al. 2018 | NCBI PRJNA523959 |
| Gastropoda | *Radix auricularia* | 910 | 4,823 | 0.58 | 2.97 | 92.7 | 87.2 | Schell et al. 2017 | NCBI PRJNA350764 |
| Gastropoda | *Chrysomallon squamiferum* | 444 | 22 | 30.20 | 49.22 | 96.6 | 87.5 | Sun et al. 2020 | NCBI PRJNA523462 |
| Gastropoda | *Achatina fulica* | 1856 | 921 | 59.59 | 116.56 | 91.5 | 95.5 | Guo et al 2019 | http://dx.doi.org/10.5524/100647 |
| Gastropoda | *Lanistes nyassanus* | 510 | 9,826 | 0.32 | 1.78 | 95.1 | 94.6 | Sun et al 2019 | NCBI PRJNA523095 |
| Gastropoda | *Marisa cornuarietis* | 535 | 665 | 4.36 | 24.07 | 96.6 | 95.1 | Sun et al 2019 | NCBI PRJNA445755 |
| Gastropoda | *Pomacea maculata* | 432 | 3,908 | 0.38 | 2.52 | 96.2 | 95.7 | Sun et al 2019 | NCBI PRJNA523958 |
| Brachiopoda | *Lingula anatina* | 406 | 2,677 | 0.46 | 2.17 | 94.5 | 97.5 | Gerdol et al. 2018 | NCBI PRJNA293030 |
| Nemertea | *Notospermus geniculatus* | 859 | 11,108 | 0.24 | 1.58 | 95.6 | 94.5 | Luo et al. 2018 | NCBI PRJNA393252 |
| Phoronida | *Phoronis australis* | 498 | 3,984 | 0.66 | 4.87 | 96.3 | 97.2 | Luo et al. 2018 | NCBI PRJNA393252 |
| Annelida | *Capitella telata* | 334 | 21,042 | 0.19 | 1.62 | 96.5 | 97.2 | Simakov et al. 2012 | NCBI PRJNA175705 |
| Annelida | *Helobdella robusta* | 235 | 1,993 | 3.06 | 13.6 | 89.6 | 92 | Simakov et al. 2012 | NCBI PFJNA259764 |

**Supplementary Table 2**: The number of repetitive elements of various types across several molluscan genomes and a brachiopod, as indicatred by RepeatModeler

| **Repetitive Element** | *Acanthopleura granulata* | *Haliotis rufescens* | *Pinctada fucata* | *Crassostrea virginica* | *Bathymodiolus platifrons* | *Scapharca broughtonii* | *Lottia gigantea* | *Lingula anatina* |
|---|---|---|---|---|---|---|---|---|
| buffer | 4 | 1 | 3 | 0 | 0 | 1 | 2 | 0 |
| DNA | 76 | 154 | 132 | 322 | 224 | 186 | 108 | 91 |
| LINE | 44 | 119 | 161 | 78 | 156 | 81 | 49 | 76 |
| SINE | 23 | 13 | 16 | 13 | 7 | 32 | 26 | 9 |
| LTR | 22 | 31 | 21 | 47 | 38 | 19 | 21 | 26 |
| RC | 10 | 13 | 0 | 85 | 52 | 60 | 3 | 6 |
| Satellite | 4 | 10 | 0 | 2 | 7 | 7 | 1 | 2 |

**Supplementary Table 3:** Comparison of heterozygosity, reproductive mode, and larval stage(s) across taxa for which short read data were available for heterozygosity analysis.

| Species | Heterozygosity | Reproductive Mode | Larval Feeding | NCBI Accession |
|---|---|---|---|---|
| *Acanthopleura granulata* | 0.653 | Broadcast spawner (Glynn 1970) | Lecithotrophic | Present study |
| *Bathymodiolus platifrons* | 0.652 | Broadcast spawner, potentially annual (Xu et al. 2018) | Planktotrophic | SRR3866232 |
| *Scapharca broughtonii* | 3.03 | Broadcast spawner, annual (Park et al. 2001, Wang et al. 2017) | Planktotrophic | SRR8535392 |
| *Crassostrea virginica* | 1.91 | Broadcast spawner, twice annually (Hayes et al. 1980) | Planktotrophic | SRR6159074 |
| *Haliotis rufescens* | 1.11 | Broadcast spawner, no clear seasonality (Giorgi et al. 1977) | Planktotrophic | SRR6761051 |
| *Pinctada fucata* | 1.03 | Broadcast spawner, annual (Behzadi 1977) | Planktotrophic | DRR001602 |
| *Lottia gigantea* | 3.15 | Broadcast spawner (Kay et al. 2005) | Lecithotrophic | SRR097541 |

**Supplementary Table 4:** Expression (TPM) of the homologs of PIF protein as identified by orthology inference. Expression is shown across the eight *A. granulata* transcriptomes generated in this study. PIF is expressed mainly in the girdle tissue.

| PIF homolog | Radula 1 (anterior) | Radula 2 | Radula 2 | Radula 4 (posterior) | Girdle | Ctenidia | Foot | Gonad |
|---|---|---|---|---|---|---|---|---|
| model.g24122.t1 | 0 | 0 | 0 | 0 | 37.28773 | 0 | 0.096919 | 0 |
| model.g24110.t1 | 0 | 0 | 0 | 0.694748 | 14.57598 | 0 | 0 | 0 |

**Supplementary Table 4:** Previously identified molluscan biomineralization genes that generated orthogroups in our analyses. For each gene, the number of orthogroups is reported, as well as the total number of gene models for each taxon across all orthogroups.

| Gene | Type | Function | Reference | Orthogroup(s) | BMOD | SBRO | PFUC | CVIR | LGIG | HRUF | AGRA | LANA |
|---|---|---|---|---|---|---|---|---|---|---|---|---|
| Ferritins | Iron | Iron binding and transport. Secreted subunits may have an immune effect. | Zhang et al 2003 | OG0002129, OG0016057 | 1 | 4 | 2 | 3 | 3 | 2 | 2 | 2 |
| Ferrochelatase | Iron | May have a role in shell pigmentation. | Song et al 2019 | OG0004417 | 1 | 1 | 1 | 2 | 1 | 1 | 1 | 1 |
| Haemocyanin | Iron | O2 courier | | OG0011679 | 0 | 0 | 1 | 1 | 0 | 3 | 1 | 1 |
| C-type Lectins | Matrix | ECM protein. Homologs known from 5 mammals and invertebrates. Known to be part of shell matrix. | Matsubara et al 2008 | OG0003537, OG00049479, OG0005252, OG0006217, OG0006279, OG0007948 | 5 | 2 | 4 | 17 | 2 | 9 | 1 | 2 |
| Calponin | Matrix | Cross links muscle fibers; may cross link proteins to stabilize shell matrix | Sleight et al 2016 | OG0004781, OG0008432, OG0014997 | 4 | 5 | 5 | 6 | 5 | 6 | 4 | 3 |
| Cartilage Matrix Protein | Matrix | Binds calcium crystals and forms part of extracellular matrix | Wang et al. 2016 | OG0001343, OG0003241, OG0006965, OG0010555 | 10 | 14 | 13 | 12 | 2 | 8 | 7 | 1 |
| Chitin Deacetylase | Matrix | Can produce chitosan from chitin. | Wang et al 2012 | OG0002391,OG0002636, OG0010102, OG0012893, OG0015720 | 5 | 8 | 5 | 5 | 5 | 6 | 9 | 3 |
| Chitin Synthase | Matrix | Synthesizes chitins, characterized by transmembrane domains in invertebrates | Weiss et al 2006 | OG0013488, OG0012243, OG0003820, OG0001957, OG0001889, OG0001819, OG0001528 | 31 | 25 | 31 | 139 | 28 | 44 | 29 | 35 |
| Chitinase | Matrix | Destruction and thus guidance of remodeling of chitin | Wang et al 2012 | OG0000213, OG0011850, OG0012263, OG0014939, OG0020247 | 12 | 12 | 11 | 6 | 9 | 17 | 4 | 8 |
| Collagens | Matrix | Structure, and may guide SMPs | Dyachuk et al 2018, Gao et al 2015, Song et al 2019 | OG000476, OG0001343, OG0003241, OG0004451, OG0004905, OG0006935, OG0010555, OG0012231, OG0013112, OG001413, OG0019628 | 4 | 6 | 4 | 9 | 7 | 4 | 3 | 6 |
| Dermatopontin | Matrix | ECM protein. Homologs known from 5 mammals and invertebrates. Known to be part of shell matrix. | Zhang et al 2006 | OG0001463 | 0 | 3 | 3 | 7 | 1 | 3 | 2 | 0 |
| Fibronectin | Matrix | Binds integrins (membrance spanning receptor proteins). Characterized from nacreous layer, mosaic pattern of loss | Marie et al 2011 | OG0000914 | 1 | 1 | 1 | 13 | 1 | 1 | 0 | 1 |
| Filamin | Matrix | Actin cross-linking | Liao et al 2015 | OG0002841 | 2 | 2 | 2 | 19 | 2 | 3 | 1 | 3 |
| Laminin | Matrix | Binds to cells and mediates attachment and organization | Aguilera et al 2017 | OG0006743, OG0007611, OG0008565, OG0016662, OG0016800, OG0019108 | 10 | 12 | 12 | 37 | 13 | 12 | 14 | 21 |
| Lustrin | Matrix | Characterized from nacreous layer; contibute to shell flexibility | Gaume et al 2014 | OG0019067 | 0 | 1 | 0 | 0 | 1 | 0 | 1 | 0 |
| Papilin | Matrix | Extracellular matrix (similar proteins exist in arthopods) glycoprotein containing several KUNTIZ domains | Gardner et al 2011 | OG0003173 | 5 | 7 | 6 | 34 | 4 | 7 | 10 | 13 |
| Perlustrin | Matrix | Similar to lustrin A. Contains growth factor receptors (GFR), but only known from Haliotis and Lottia. | Weiss et al 2000 | OG0014184 | 0 | 2 | 1 | 0 | 0 | 1 | 0 | 0 |
| Vitellogenin | Matrix | lipid transporter | | OG0011518 | 2 | 1 | 1 | 11 | 0 | 1 | 1 | 1 |
| BMSP | Mineralization | Binds calcium carbonate crystals. Can bind calcite and aragonite. Is preprotein that breaks into BMSP120 (4 VWAs, 1 chitin-binding domain) and BMSP100 (phosporylated and part of nacreous layer, dense in myostracum). Homologous to PIF177 | Suzki et al 2011 | OG0003624, OG0015921 | 1 | 3 | 2 | 1 | 1 | 1 | 2 | 0 |
| Calcitonin Receptor | Mineralization | GPCR that binds calcitonin (peptide hormone) and maintains calcium homeostasis | | OG0001242, OG0002433, OG0001719 | 3 | 13 | 10 | 13 | 6 | 6 | 7 | 5 |
| Follistatin | Mineralization | Chitin binding, protein interactions | Feng et al 2017 | OG0010890, OG0001081 | 1 | 1 | 1 | 1 | 1 | 1 | 1 | 1 |
| Nacrein | Mineralization | Synthesizes HCO3- | Miyamoto et al 2005 | OG0001347 | 0 | 0 | 0 | 0 | 1 | 1 | 2 | 2 |
| Perlucin | Mineralization | Increases precipitation of calcium carbonate. Contains C-type lectin domain. | Weiss et al 2000 | OG0004979, OG0006217 | 3 | 5 | 4 | 15 | 4 | 7 | 1 | 1 |
| Peroxidases | Mineralization | May stimulate collagen production | Feng et al 2017, DeNichilo et al 2016 | OG0000097,OG0001338, OG0003933,OG0006494,OG0011251 | 14 | 15 | 13 | 21 | 4 | 17 | 13 | 8 |
| PIF80/97 Precursor | Mineralization | Binds to chitin, implicated in crystal growth patterns | Suzuki et al 2013 | OG0000555, OG0003624 | 3 | 7 | 7 | 6 | 6 | 1 | 4 | 0 |
| Secreted protein acidic and rich in Cysteine (SPARC) | Mineralization | Calciium binding, stabilizes vaterite to inhibit calcite formation (can facilitate nacre formation) | Xie et al 2016 | OG0006587 | 1 | 1 | 1 | 3 | 1 | 0 | 1 | 1 |
| Tyrosinase | Mineralization | Crosslinks o-diphenols and aids in formation of the periostracal layer | Mann et al 2012 | OG0000131 | 16 | 18 | 6 | 13 | 1 | 1 | 3 | 7 |
| Carbonic Anhydrases | Mineralization | Interconversion of water and CO2 to carbonic acid, protons, and bicarbonate ions. | Duvail et al 1997 | OG0000554,OG0001347,OG0001347,OG0015230 | 10 | 10 | 9 | 17 | 8 | 10 | 4 | 6 |
| Activator protein-1 (AP-1) | Trans. Factor | Binds directly to KRMP and enhances promoter activity | Zheng et al 2015 | OG0008602 | 1 | 1 | 1 | 1 | 1 | 1 | 1 | 1 |
| protein (ACCBP) | Trans. Factor | grows of aragonite crystal faces to facilitate nacre development | Ma et al 2007 | OG0014423, OG0011654, OG0015373 | 22 | 27 | 18 | 40 | 20 | 24 | 15 | 16 |
| PfMSX | Trans. Factor | Binds directly to Pearlin and enhances promoter activity | Zhao et al 2014 | OG0001643 | 50 | 47 | 36 | 57 | 19 | 36 | 47 | 47 |
| Pf-POU3F4 | Trans. Factor | | Gao et al 2016 | OG0018960 | 4 | 4 | 3 | 7 | 4 | 6 | 4 | 5 |
| Rel | Trans. Factor | Binds directly to Nacrein and enhances promoter activity | Sun et al 2015 | OG0003590, OG0011250 | 2 | 2 | 2 | 3 | 2 | 2 | 3 | 2 |

**Supplementary Table 6:** Total number of gene models in molluscan genome annotations before and after clustering with CDHit, and the number of high-quality IRE predictions from SIRE for each set of clustered gene models across molluscan taxa.

| Taxon | Initial Gene | Clustered Gene | High-quality | NCBI Project Accession or Other |
|---|---|---|---|---|
| *Acanthopleura granulata* | 81691 | 20470 | 271 | PRJNA578131 |
| *Bathymodiolus platifrons* | 33584 | 27449 | 181 | PRJNA328542 |
| *Crassostrea virginica* | 66625 | 35117 | 138 | PRJNA376014 |
| *Haliotis rufescens* | 57785 | 48537 | 72 | https://abalone.dbgenome.org/ |
| *Lottia gigantea* | 23851 | 21739 | 126 | PRJNA175706 |
| *Pinctada fucata* | 31477 | 29486 | 137 | https://marinegenomics.oist.jp |
| *Scapharca broughtonii* | 24045 | 22288 | 201 | PRJNA193911 |

**Supplementary Table 5:** Translated gene models identified by SilkSlider as putative genes for proteins with silk-like domains. A top blastp hit is listed for each protein, along with the name/gene/accession of the most similar result and E-value of the blastp hit. The final column is a separate analysis via SignalP-5.0 of whether a signal peptide is predicted for this gene model (Likelihood % in parentheses).

| Gene model | Top blastp hit | Taxon | Accession | Evalue | Signal Peptide Predicted |
|---|---|---|---|---|---|
| >evm.model.scf7180000027668.320785.4\|m.11338 | carbonic anhydrase 14-like | *Crassostrea virginica* | XP_022339696.1 | 6.00E-56 | Yes, 0.9555 |
| >model.g30149.t1\|m.35336 | chitinase (acidic mammalian) | *Mizuhopecten yessoensis* | OWF47140.1 | 5.00E-06 | Yes, 0.9797 |
| >evm.model.Super-Scaffold_100008.944\|m.4245 | collagen alpha-2 chain-like | *Acanthaster planci* | XP_022105828.1 | 6.00E-96 | Yes, 0.7671 |
| >model.g17066.t1\|m.32742 | collagen alpha-2 chain-like | *Mizuhopecten yessoensis* | XP_021346726.1 | 0.00E+00 | Yes, 0.9980 |
| >model.g4747.t1\|m.29327 | complement C1g subcomponent B-like | *Octopus vulgaris* | XP_029636373.1 | 3.00E-94 | Yes, 0.8324 |
| >model.g9371.t1\|m.16792 | deodyribonuclease-1-like | *Mizuhopecten yessoensis* | XP_021351957.1 | 2.00E-59 | Yes, 0.7191 |
| >model.g6261.t1\|m.14920 | eukaryotic translation initiation factor 3 subunit A-like | *Octopus bimaculoides* | XP_014778366.1 | 4.00E-26 | Yes, 0.7402 |
| >evm.model.Super-Scaffold_100004.800\|m.2401 | fibril-forming collagen alpha chain-like | *Octopus vulgaris* | XP_029638258.1 | 0.00E+00 | Yes, 0.9996 |
| >model.g6411.t1\|m.15452 | fibrillar collagen (many annelid hits) | *Riftia pachyptila* | AAF80453.1 | 2.00E-57 | Yes, 0.9994 |
| >model.g10834.t1\|m.17317 | frizzled-5-like, membrance frizzled | *Crassostrea gigas* | XP_011453204.1 | 1.00E-69 | Yes, 0.9714 |
| >evm.model.Super-Scaffold_100003.741\|m.963 | Histone-lysine N-methyltransferase PRDM9 | *Mizuhopecten yessoensis* | OWF46683.1 | 7.00E-14 | No |
| >model.g5858.t1.1.5d3b846d\|m.29547 | IgGFc-binding protein-like | *Crassostrea gigas* | XP_011418167.1 | 1.00E-105 | Yes, 0.9833 |
| >model.g18955.t1\|m.22080 | myosin-IIIb isoform X2 | *Lingula anatina* | XP_013418100.1 | 0.00E+00 | No |
| >model.g4723.t1\|m.14290 | nucleolin-like | *Lingula anatina* | XP_013398714.2 | 5.00E-28 | Yes, 0.6206 |
| >model.g25880.t1\|m.25261 | protein SSXT-like | *Mizuhopecten yessoensis* | XP_021354255.1 | 2.00E-30 | No |
| >model.g29156.t1\|m.26325 | sialic acid binding lectin | *Helix pomatia* | ABF00124.1 | 6.00E-09 | Yes, 0.9978 |
| >model.g25029.t1\|m.24855 | uncharacterized | *Lingula anatina* | XP_013408830.1 | 4.00E-92 | Yes, 0.9609 |
| >model.g6163.t1\|m.29750 | ptotocadherin Fat 4-like | *Lingula anatina* | XP_013414156.1 | 9.00E-131 | Yes, 0.9933 |
| >model.g10732.t1\|m.17549 | uncharacterized from several molluscs | *Crassostrea gigas* | XP_011429067.1 | 2.00E-12 | Yes, 0.7249 |
| >evm.model.Super-Scaffold_100007.378\|m.3990 | uncharacterized PE-PGRS family protein PE | *Diabrotica virgifera* | XP_028143928.1 | 4.50E+00 | Yes, 0.9988 |
| >model.g11190.t1.1.5d3b864d\|m.31279 | uncharacterized, predicted flocculation (FLO10),mucin5 | *Crassostrea gigas* | XP_011424285.1 | 9.00E-62 | Yes, 0.9103 |
| >evm.model.Super-Scaffold_100011.870\|m.5327 | no hits | | | | Yes, 0.9849 |
| >evm.model.Super-Scaffold_158.18\|m.10151 | no hits | | | | Yes, 0.9761 |
| >model.g12987.t1\|m.18861 | no hits | | | | Yes, 0.9938 |
| >model.g14042.t1\|m.19238 | no hits | | | | Yes, 0.6893 |
| >model.g16712.t1\|m.21018 | no hits | | | | Yes, 0.8798 |
| >model.g28544.t1\|m.26272 | no hits | | | | Yes, 0.9868 |
| >model.g28545.t1\|m.35013 | no hits | | | | Yes, 0.7622 |
| >model.g289.t1\|m.11469 | no hits | | | | Yes, 0.9815 |
| >model.g31005.t1\|m.27320 | no hits | | | | No |
| >evm.model.Super-Scaffold_100011.762\|m.5377 | no hits | | | | Yes, 0.9959 |

**Supplementary Table 7:** Query sequences used for each biomineralization gene of interest, indicating the species the sequence is from and the NCBI accession number of each.

| Biomineralization Gene | Query Species | Accession |
|---|---|---|
| Activator protein-1 (AP-1) | *Haliotis discus discus* | ADQ43242.1 |
| Amorphous calcium carbonate binding protein (ACCBP) | *Pinctada fucata* | ABF13208.1 |
| BMSP | *Pinctada fucata* | AYN73066.1 |
| Calcitonin Receptor | *Crassostrea gigas* | QDH43372.1 |
| Calponin | *Mizuhopecten yessoensis* | BAP84555.1 |
| Carbonic Anhydrases | *Mizuhopecten yessoensis* | OWF36341.1 |
| Cartilage Matrix Protein | *Pinctada fucata* | AQN80778.1 |
| Chitin Deacetylase | *Hyriopsis cumingii* | AFO53262.1 |
| Chitin Synthase | *Pinctada fucata* | BAF73720.1 |
| Chitinase | *Octopus vulgaris* | X02571.1 |
| Collagens | *Haliotis rufescens* | AIZ03373.1 |
| C-type Lectins | *Pinctada fucata* | ADX95743.1 |
| Dermatopontin | *Pinctada imbricata* | AFK64754.1 |
| Ferritins | *Mizuhopecten yessoensis]* | AHH31563.1 |
| Ferrochelatase | *Pomacea canaliculata* | XP_025082221.1 |
| Fibronectin | *Mizuhopecten yessoensis* | OWF41565.1 |
| Filamin | *Euprymna scolopes* | AAT99400.1 |
| Follistatin | *Mizuhopecten yessoensis* | OWF54001.1 |
| Haemocyanin | *Aplysia californica* | CAD88977.1 |
| Laminin | *Crassostrea virginica* | XP_022314761.1 |
| Lustrin | *Haliotis tuberculata* | ADM52208.2 |
| Nacrein | *Pinctada fucata* | BAA11940.1 |
| Papilin | *Mizuhopecten yessoensis* | OWF36203.1 |
| Perlucin | *Mizuhopecten yessoensis* | OWF55388.1 |
| Perlustrin | *Mizuhopecten yessoensis* | OWF44587.1 |
| Peroxidases | *Pinctada fucata* | ALK82327.1 |
| Pf-POU3F4 | *Pinctada fucata* | AKJ32469.1 |
| PIF80/97 Precursor | *Pinctada margaritifera* | BAM66823.1 |
| Rel | *Mizuhopecten yessoensis* | AKC01670.1 |
| Secreted protein acidic and rich in Cysteine (SPARC) | *Pinctada fucata* | AND99565.1 |
| Tyrosinase | *Mizuhopecten yessoensis* | AKE79095.1 |
| Vitellogenin | *Scapharca broughtonii* | AYE92811.1 |

Supplementary Document 1: All programs, configuration files, and code used in this study.

| Program | Function in present study | Version |
|---|---|---|
| Alignment_Compare | Removing sequences that don't meet minimum overlap requirements within orthogroups | 1.0 |
| Augustus | Gene prediction | 3.3.2 |
| BEDTools | Combining multiple sets of gene predictions | 2.29.2 |
| Bionano Solve | Using optical mapping data to scaffold | 3.4 |
| Blast+ | Identifying orthogroups that matched biomineralization genes | 2.10.0 |
| BlobTools | Assessing genome contamination, and generation of snail plot | 2.0 |
| BMGE | Trim ambiguously aligned columns from orthogroup alignments | 1.12 |
| Bowtie | Mapping PE data back to genome in preparation for PILON | 2.2.5 |
| BUSCO | Assessing genome completeness and duplication | 4.0.2 / 3.9 |
| CDHit | Markov clustering of transcripts and gene models | 4.8.1 |
| EdgeR | Generating TPM values for expression data | 3.28.1 |
| EVM | Tools within used for sequence analysis | 1.1.1 |
| FastTree | Initial trees within orthogroups to give PhyloPyPruner | 2.1 |
| HMMCleaner | Remove sequence regions that are misaligned in orthogroups | 0.180750 |
| IQTree | Phylogenetic tree building | 1.6.12 |
| MAFFT | Aligning sequences for phylogenetic tree building | 7.45 |
| Maker | Annotating genome to generate training files for SNAP and Augustus | 2.31.10 |

| | | |
|---|---|---|
| MaSuRCA | Hybrid assembly of genome from paired-end and long-read sequencing data | 3.3.5 |
| OrthoFinder | Determination of orthogroups between taxa | 2.3.8 |
| PASA | Mapping transcriptomic data back to genome assembly to generate an annotation | 2.3.3 |
| PhyloPyPruner | Prune paralogs from orthology inferences after OrthoFinder | 1.0 |
| PILON | Map paired-end data back to genome assembly and correct potential errors produced by long read sequencing | 1.23 |
| PoreChop | Trim nanopore-specific adapter sequences, and discard chimeric sequences | 0.2.4 |
| QUAST | Assess genome quality | 5.0.2 |
| RaxML | Generate maximum likelihood phylogenies | 8.2.12 |
| Reapr | Map paired end data back to final genome assembly to assess completeness | 1.0.18 |
| Redundans | Collapse contigs that differ due to heterozygosity in genome assembly | 0.14a |
| RepeatMasker | Generate a file to soft-mask repetitive regions in the genome during annotation | 4.0.9 |
| RepeatModeler | Determine the repetitive content of genomes | 2.0 |
| Salmon | Map transcriptome reads back to genome and generate quantifications per isoform | 0.11.3 |
| SilkSlider | Identify possible silk-like proteins | 0.2.2 |
| SIRE | Identify potential IREs | 2.0 |
| SNAP | *De novo* gene prediction | 5th update |
| SynMap (COGE) | Generate dot plots to represent synteny between genomes | 2.0 |

| | | |
|---|---|---|
| Transdecoder | Identify the longest open reading frame for each gene model and translate to protein sequence | 5.5.0 |
| Trinity | Assembly transcriptomes | 2.8.4 |
| uniqHaplo.pl | Removing redundant sequences from orthogroups | 0.1.4 |

### Genome Assembly:

#### MaSuRCA:

masurcaconfig.txt file:

```
DATA
#HiSeq X with 2 X 150 bp reads
PE= pe 150 20 AGRA_1.fastq AGRA_2.fastq
NANOPORE=nanoporecombined.fastq
END

PARAMETERS
EXTEND_JUMP_READS=0
GRAPH_KMER_SIZE = auto
USE_LINKING_MATES =1
LIMIT_JUMP_COVERAGE = 300
CA_PARAMETERS = cgwErrorRate=0.15
KMER_COUNT_THRESHOLD = 1
CLOSE_GAPS=1
NUM_THREADS = 16
JF_SIZE = 200000000
SOAP_ASSEMBLY=0
END
```

#### Quast to evaluate genome assemblies:

```
use quast
quast.py -t 8 --plots-format pdf genome.fasta
```

#### TrimGalore to trim PE data before mapping:

```
use trim_galore
mkdir trimmed
trim_galore -fastqc --paired AGRA_1.fastq AGRA_2.fastq -o trimmed
```

#### Bowtie mapping:

```
bowtie2 -p 8 -x scaffolds_bt2_index -1 AGRA_1.fq -2 AGRA_2.fq |
samtools view -Sb - > PE_mapped_for_pilon.corrected.sam
samtools view -bh PE_mapped_for_pilon.corrected.sam -o
PE_mapped_for_pilon.corrected.bam
samtools sort PE_mapped_for_pilon.corrected.bam -o
PE_mapped_for_pilon.corrected.sorted.bam
samtools index PE_mapped_for_pilon.corrected.sorted.bam
samtools depth PE_mapped_for_pilon.corrected.sorted.bam > depth.txt
```

#### PILON:

```
srun java -Xmx256G -jar /share/apps/bioinfoJava/pilon-1.23.jar --
genome scaffolds.fasta --bam PE_mapped_for_pilon.corrected.sorted.bam
--output AGRA.pilon1.corrected --outdir PILON_output_1 --changes --
diploid  --verbose
```

**Redundans:**

```
use redundans
redundans.py -t 16 -m 255 --noscaffolding --norearrangements -i
AGRA_1.fastq AGRA_2.fastq -f final.genome.scf.fasta
```

### Genome Annotation:

**Overall Approach:**
Maker was run twice with transcriptome evidence as est2genome and multiple other chitons' protein evidence. Each time, the best-scoring set of proteins +1000 framing basepairs were removed into new files, and used to train Augustus. SNAP was also trained each cycle. The training file produced by the second run was carried forward into Augustus *de novo* predictions. The trained Augustus performed much better (when species=fly BUSCO obd9 complete was 32%, versus with species=agra_round2, 92%).

PASA was run to map transcripts back to the repeat-masked genome, using the composite transcriptome cleaned according to PASA built-in process.

EVM would not work on chitons. Examination of outputs seems to indicate that chitons often alternatively splice exon 1, and EVM tended to discard predictions where this discordance existed.

PASA and Augustus outputs were combined, and bedtools intersect used to remove perfectly redundant models. This left the first set of 81K gene models. CDHit was run to cluster these based on probably redundancy, and at a similarity cutoff of 0.8 produced the 20K gene model set.

**Repeat Modeling and Masking:**
```
RepeatModeler/BuildDatabase -name AGRA -engine ncbi
/home/scutopus/maker/2018_12_20_AGRA_post_redundans_optical/AGRAredund
ansoptical.fasta
RepeatModeler/RepeatModeler -pa 4 -engine ncbi -database AGRA 2>&1 |
tee repeatmodeler.log
```

**Maker Run 1:**
Thanks to maker resources at:
http://weatherby.genetics.utah.edu/MAKER/wiki/index.php/The_MAKER_control_files_explained
https://github.com/sujaikumar/assemblage/blob/master/README-annotation.md
https://gist.github.com/darencard/bb1001ac1532dd4225b030cf0cd61ce2

Run to generate a first pass set of gene models. Note est2genome=1 using composite *A. granulata* transcriptome for most complete annotation possible. Protein evidence incorporated from several additional chiton species as listed.

Maker1_opts.ctl file:

```
#-----Genome (these are always required)
##Added genome file here:
genome=scaffolds.fasta
```

```
organism_type=eukaryotic #eukaryotic or prokaryotic. Default is
eukaryotic

#-----Re-annotation Using MAKER Derived GFF3
maker_gff= #MAKER derived GFF3 file
est_pass=0 #use ESTs in maker_gff: 1 = yes, 0 = no
altest_pass=0 #use alternate organism ESTs in maker_gff: 1 = yes, 0 =
no
protein_pass=0 #use protein alignments in maker_gff: 1 = yes, 0 = no
rm_pass=0 #use repeats in maker_gff: 1 = yes, 0 = no
model_pass=0 #use gene models in maker_gff: 1 = yes, 0 = no
pred_pass=0 #use ab-initio predictions in maker_gff: 1 = yes, 0 = no
other_pass=0 #passthrough anything else in maker_gff: 1 = yes, 0 = no

#-----EST Evidence (for best results provide a file for at least one)
est=../Acanthopleura_granulata_KK701_composite_cd-hit.fas #set of ESTs
or assembled mRNA-seq in fasta format
altest= #EST/cDNA sequence file in fasta format from an alternate
organism
est_gff= #aligned ESTs or mRNA-seq from an external GFF3 file
altest_gff= #aligned ESTs from a closely related species in GFF3
format

#-----Protein Homology Evidence (for best results provide a file for
at least one)
protein= Acanthochitona_crinita-
SRR5110525.fa.transdecoder.pep,Acanthopleura_gemmata_contaminant_free_
trinity.fa.transdecoder.pep,Acanthopleura_granulata_KK701_composite_cd
-hit.fa.transdecoder.pep
Callochiton_sp-
Pl49_4C.fa.transdecoder.pep,Chaetopleura_apiculata_girdle.fa.transdeco
der.pep,Chiton_marmoratus-
Speiser.fa.transdecoder.pep,Hanleya_nagelfar_mantle.fa.transdecoder.pe
p,Leptochiton_asellus-
mantle.fa.transdecoder.pep,Mopalia_muscosa_KK364-1-
4R.fa.transdecoder.pep,Tonicella_lineata_KK374-1-
4R.fa.transdecoder.pep #protein sequence file in fasta format (i.e.
from multiple organisms)
protein_gff=  #aligned protein homology evidence from an external GFF3
file

#-----Repeat Masking (leave values blank to skip repeat masking)
model_org=all #select a model organism for RepBase masking in
RepeatMasker
##AGRA repeat library added!
```

```
rmlib=../consensi.fa #provide an organism specific repeat library in
fasta format for RepeatMasker
repeat_protein=/share/apps/maker/data/te_proteins.fasta #provide a
fasta file of transposable element proteins for RepeatRunner
rm_gff= #pre-identified repeat elements from an external GFF3 file
prok_rm=0 #forces MAKER to Repeatmask prokaryotes (no reason to change
this), 1 = yes, 0 = no
softmask=1 #use soft-masking rather than hard-masking in BLAST (i.e.
seg and dust filtering)

#-----Gene Prediction
snaphmm= #SNAP HMM file#SNAP HMM file
gmhmm= #GeneMark HMM file
augustus_species=fly
fgenesh_par_file= #FGENESH parameter file
pred_gff= #ab-initio predictions from an external GFF3 file
model_gff= #annotated gene models from an external GFF3 file
(annotation pass-through)
## Changed to 1 to allow transcriptome to be used directly to annotate
est2genome=1 #infer gene predictions directly from ESTs, 1 = yes, 0 =
no
protein2genome=1 #infer predictions from protein homology, 1 = yes, 0
= no
trna=0 #find tRNAs with tRNAscan, 1 = yes, 0 = no
snoscan_rrna= #rRNA file to have Snoscan find snoRNAs
unmask=1 #also run ab-initio prediction programs on unmasked sequence,
1 = yes, 0 = no

#-----Other Annotation Feature Types (features MAKER doesn't
recognize)
other_gff= #extra features to pass-through to final MAKER generated
GFF3 file

#-----External Application Behavior Options
alt_peptide=C #amino acid used to replace non-standard amino acids in
BLAST databases
cpus=1 #max number of cpus to use in BLAST and RepeatMasker (not for
MPI, leave 1 when using MPI)

#-----MAKER Behavior Options
max_dna_len=100000 #length for dividing up contigs into chunks
(increases/decreases memory usage)
min_contig=1 #skip genome contigs below this length (under 10kb are
often useless)
```

```
pred_flank=200 #flank for extending evidence clusters sent to gene
predictors
pred_stats=0 #report AED and QI statistics for all predictions as well
as models
AED_threshold=1 #Maximum Annotation Edit Distance allowed (bound by 0
and 1)
min_protein=0 #require at least this many amino acids in predicted
proteins
alt_splice=0 #Take extra steps to try and find alternative splicing, 1
= yes, 0 = no
always_complete=0 #extra steps to force start and stop codons, 1 =
yes, 0 = no
map_forward=0 #map names and attributes forward from old GFF3 genes, 1
= yes, 0 = no
keep_preds=0 #Concordance threshold to add unsupported gene prediction
(bound by 0 and 1)

split_hit=10000 #length for the splitting of hits (expected max intron
size for evidence alignments)
single_exon=1 #consider single exon EST evidence when generating
annotations, 1 = yes, 0 = no
single_length=250 #min length required for single exon ESTs if
'single_exon is enabled'
correct_est_fusion=0 #limits use of ESTs in annotation to avoid fusion
genes

tries=2 #number of times to try a contig if there is a failure for
some reason
clean_try=0 #remove all data from previous run before retrying, 1 =
yes, 0 = no
clean_up=0 #removes theVoid directory with individual analysis files,
1 = yes, 0 = no
TMP= #specify a directory other than the system default temporary
directory for temporary files
```

**Reformat outputs for training:**

```
gff3_merge -s -d maker1_master_datastore_index.log >
maker1.all.maker.gff
fasta_merge -d maker1_master_datastore_index.log
gff3_merge -n -s -d maker1_master_datastore_index.log >
maker1.all.maker.noseq.gff
```

**SNAP training:**

```
mkdir snap/round2
```

Using an AED cutoff of 0.25 to pull the higher quality models:

```
maker2zff -x 0.25 -l 50 -d ../../maker2_master_datastore_index.log
```
This produces a .ann and a .dna file.
Everything must be renamed carefully or SNAP will fail to find the files later:
```
rename 's/genome/AGRA_rnd2.zff.length50_aed0.25/g' *
```
We now pull these gene models with 1000 basepairs of extra sequence to make sure we get the entire gene:
```
fathom AGRA_rnd2.zff.length50_aed0.25.ann
AGRA_rnd2.zff.length50_aed0.25.dna -gene-stats > gene-stats.log 2>&1
fathom AGRA_rnd2.zff.length50_aed0.25.ann
AGRA_rnd2.zff.length50_aed0.25.dna -validate  > validate.log 2>&1
fathom AGRA_rnd2.zff.length50_aed0.25.ann
AGRA_rnd2.zff.length50_aed0.25.dna -categorize 100 > categorize.log
2>&1
fathom uni.ann uni.dna -export 1000 -plus > uni-plus.log 2>&1
```
Now make a directory for SNAP to find:
```
mkdir params
cd params
forge ../export.ann ../export.dna > ../forge.log 2>&1
cd ..
hmm-assembler.pl AGRA_rnd2.zff.length50_aed0.25 params >
AGRA_rnd2.zff.length50_aed0.25.hmm
```

**Augustus Training**
```
mkdir augustus
cd augustus
awk -v OFS="\t" '{ if ($3 == "mRNA") print $1, $4, $5 }'
../maker2_rnd2.all.maker.noseq.gff | \
  awk -v OFS="\t" '{ if ($2 < 1000) print $1, "0", $3+1000; else print
$1, $2-1000, $3+1000 }' | \
  bedtools getfasta -fi ../../AGRAredundansoptical.fasta -bed - -fo
maker2_rnd2.all.maker.transcripts1000.fasta
```
We will use Augustus within BUSCO to create the training file:
```
BUSCO.py -i maker2_rnd2.all.maker.transcripts1000.fasta  -o
maker2_rnd2_maker -l /usr/bin/busco/metazoa_odb9/ \
  -m genome -c 8 --long -sp fly -z --augustus_parameters='--
progress=true'
```
We also get a glimpse of our “completeness” - here we had C91.5%
[S:90.1%,D:1.4%],F:2.6%,M:5.9%,n:978
This produces a directory 'run_maker2_rnd2_maker/augustus_output/retraining_parameters
Now we rename the output files. ***This re-naming step is vital, or Augustus won’t find anything:
```
rename 's/BUSCO_maker2_rnd2_maker_2594518820/agra_round2/g' *
sed -i 's/BUSCO_maker2_rnd2_maker_2594518820/agra_round2/g'
agra_round2_parameters.cfg
sed -i 's/BUSCO_maker2_rnd2_maker_2594518820/agra_round2/g'
agra_round2_parameters.cfg.orig1
```

Now these need to be copied into the AUGUSTUS HMM directory so they can be located by Augustus. Make sure you copy to the version of Augustus called by maker specifically (check the maker_exe.ctl file).
Make a new directory within augustus/config/species.
*** The name of this directory MUST MATCH the training files. These training files were "agra_round2"

```
cp agra_round2* /home/.../augustus-3.2.2/config/species/agra_round2
```

In subsequent augustus runs, SPECIES=agra_round2

Count the number of gene models after each round of maker:

```
cd maker1.maker.output
cat maker1.all.maker.gff | awk '{ if ($3 == "gene") print $0 }' | awk
'{ sum += ($5 - $4) } END { print NR, sum / NR }'
```

And view a histogram of AED scores to mark improvement:

```
perl /home/scutopus/maker/bin/AED_cdf_generator.pl -b 0.025
maker1.all.maker.gff
```

Output: 39488 6450.59

**Maker Run 2**:
Used the same parameters as run 1, but incorporated the training files for both SNAP and Augustus as listed above.

**Running trained Augustus on its own:**

```
augustus --gff3=on --outfile=augustus.gff3 --species=agra_round2
scaffolds.fasta
```

**PASA Run 1:**

```
/share/apps/pasa/PASApipeline-v2.3.3/Launch_PASA_pipeline.pl \
    -c alignassembly.config -C -R -g ../4addedscaffolds.fasta \
    -t Acanthopleura_granulata_KK701_composite_cd-hit.fasta.clean -T -
u Acanthopleura_granulata_KK701_composite_cd-hit.fasta \
    --ALIGNERS blat,gmap --CPU 1
### Changed CPU to 1 because of known issues using SQLite and
multithreading
```

**Using EVM to move from .gff3 to CDS:**

```
use evm
srun --mpi=pmi2 -n 1  *.gff3 ../4addedscaffolds.fasta CDS > CDS.fasta
```

**Combining features from multiple annotations:**

```
bedtools merge -i *.gff3
```

**Transdecoder:**

```
#!/bin/bash
for FILENAME in *.fasta
do
TransDecoder.LongOrfs -t $FILENAME
done

for FILENAME in ./*/longest_orfs.pep
do
DIR=`echo $FILENAME | cut -d "/" -f 1-2`
echo $FILENAME
echo $DIR
blastp -query $FILENAME -db
/home/wirenia/blast_dbs/uniprot_sprot_21_Sept_2016/uniprot_sprot.fasta
-max_target_seqs 1 -outfmt 6 -evalue 1e-5 -num_threads 30 -out
$DIR/longest_orfs.pep.blast_results
done

for FILENAME in ./*.transdecoder_dir/longest_orfs.pep
do
hmmscan --cpu 30 --domtblout $FILENAME.domtblout
/home/wirenia/blast_dbs/Pfam27/Pfam-A.hmm $FILENAME
done

for FILENAME in *.fa
do
TransDecoder.Predict --cpu 30 -t $FILENAME --retain_long_orfs --
single_best_orf --retain_pfam_hits
./$FILENAME.transdecoder_dir/longest_orfs.pep.domtblout --
retain_blastp_hits
./$FILENAME.transdecoder_dir/longest_orfs.pep.blast_results
done
```

### Genomic Comparisons:

**RepeatModeler:**
Run on all downloaded mollusc genomes for comparisons:

```
use repeatmodeler-1.0.8

BuildDatabase -name *.seqfile -engine ncbi *-scaffolds.fas
RepeatModeler -database *.seqfile >& *.seqfile.out
```

**Reapr:**
Using Reapr on trimmed PE data to determine genome completeness:

```
reapr smaltmap -n 2 $GENOME $PE_1 $PE2 $GENOME.mapped.bam
reapr perfectmap reapr_renamed_4added.fa AGRA_1.fastq AGRA_2.fastq 150
perfect
```

**GenomeScope:**
Obtaining reads.histo files from jellyfish for other molluscan genomes for GenomeScope:

```
use trinity
me=`whoami`
fastq-dump --defline-seq '@$sn[_$rn]/$ri' --split-files SRR*******
rm -rf /home/$me/ncbi/
srun jellyfish count -C -m 21 -s 100M -t 12 *.fq -o reads.jf
srun jellyfish histo -t 12 reads.jf > reads.histo
```

**Salmon:**
Salmon mapping transcriptome(s) back to genome for quantification files to generate expression data:

```
use salmon
#index genome-based transcripts
salmon index -t Acanthopleura_granulata-transcripts.fas -i
transcripts_index -k 31
#Quantify transcriptomes
salmon quant -i transcripts_index -l A -1 KK701-*R_1.fastq.gz -2
KK701-*R_2.fastq.gz --validateMappings -o KK701.*.transcripts_quant
```

**CDHit and CDHit-EST:**

```
cdhit -i Combined.fasta -o cdhitCombined.fas -n 4 -c 0.8 -T 10
cdhit-est -i $file -o $file.0.8.cdhit -c 0.8 -T 10
```

**SIRE:**
Uploaded each taxon’s clustered file to SIRE2.0 WebServer

**Ortholog Searching:**

```
orthofinder -t 20 -I 2.1 -M msa -T fasttree -f ./pep -o
./OrthoFinder_results
```

**OrthoFinder CleanUp:**

```
#Delete sequences shorter than $MIN_SEQUENCE_LENGTH
echo "Deleting sequences shorter than $MIN_SEQUENCE_LENGTH AAs..."
for FILENAME in *.fa
do
grep -B 1 "[^>].\{$MIN_SEQUENCE_LENGTH,\}" $FILENAME > $FILENAME.out
sed -i 's/--//g' $FILENAME.out
sed -i '/^$/d' $FILENAME.out
rm -rf $FILENAME
mv $FILENAME.out $FILENAME
done
echo Done

#If fewer than $MIN_TAXA different species are represented in the
file, move that file to a "rejected_few_taxa" directory.
echo "Removing groups with fewer than $MIN_TAXA taxa..."
mkdir -p rejected_few_taxa_1
for FILENAME in *.fa
do
awk -F"|" '/^>/{ taxon[$1]++ } END{for(o in taxon){print o,taxon[o]}}'
$FILENAME > $FILENAME\.taxon_count #Creates temporary file with taxon
abbreviation and number of sequences for that taxon in $FILENAME
taxon_count=`grep -v 0 $FILENAME\.taxon_count | wc -l` #Counts the
number of lines with an integer >0 (= the number of taxa with at least
1 sequence)
if [ "$taxon_count" -lt "$MIN_TAXA" ] ; then
echo $FILENAME
mv $FILENAME ./rejected_few_taxa_1/
fi
done
rm -rf *[0-9].fa.taxon_count
echo Done
echo


#Remove redundant sequences using uniqHaplo
(http://doi.org/10.5281/zenodo.166024)
mkdir preUniqHaplo
cp *.fa preUniqHaplo
echo "Removing redundant sequences using uniqHaplo..."
```

```
ls *[0-9].fa | parallel -j $CORES 'perl /usr/bin/uniqHaplo.pl -a {} >
{}.uniq'
rm -rf *.fa
rename 's/.fa.uniq/.fa/g' *.fa.uniq
echo Done
echo

#Align the remaining sequences using Mafft.
echo "Aligning sequences using Mafft (auto)..."
mkdir backup_alignments
ls *[0-9].fa | parallel -j $CORES 'mafft --auto --localpair --
maxiterate 1000 {} > {}.aln'
rm -rf *[0-9].fa
rename 's/.fa.aln/.fa/g' *.fa.aln
cp *.fa ./backup_alignments/
echo Done
echo

#Remove newlines.
echo "Removing linebreaks in sequences..."
for FILENAME in *.fa
do
sed -i ':a; $!N; /^>/!s/\n\([^>]\)/\1/; ta; P; D' $FILENAME
done
echo Done
echo

#Clean alignments with HmmCleaner
echo "Removing misaligned sequence regions with HmmCleaner..."
mkdir HmmCleaner_files
ls *.fa | parallel -j $CORES 'HmmCleaner.pl {} --specificity'
mv *.fa ./HmmCleaner_files
mv *.log ./HmmCleaner_files
mv *.score ./HmmCleaner_files
rename 's/_hmm.fasta/.fa/g' *_hmm.fasta
echo Done
echo


#Trim alignments with BMGE
echo "Trimming ambiguously aligned columns in the alignment with
BMGE..."
mkdir backup_pre-BMGE
ls *.fa | parallel -j $CORES 'java -jar /usr/bin/BMGE-1.12/BMGE.jar -i
{} -t AA -of {}.BMGE'
```

```
mv *.fa ./backup_pre-BMGE
cp *.BMGE ./backup_pre-BMGE
rename 's/.BMGE//g' *.BMGE
echo Done
echo

#Remove newlines.
echo "Removing linebreaks in sequences..."
for FILENAME in *.fa
do
sed -i ':a; $!N; /^>/!s/\n\([^>]\)/\1/; ta; P; D' $FILENAME
done
echo Done
echo

#Remove any sequences that don't overlap with all other sequences by
at least 20 amino acids.
for FILENAME in *.fa
do
java -cp /usr/bin AlignmentCompare $FILENAME
done
echo Done
echo
rm -rf myTempFile.txt

#If fewer than $MIN_TAXA different species are represented in the
file, move that file to the "rejected_few_taxa" directory.
echo "Removing groups with fewer than $MIN_TAXA taxa..."
mkdir -p rejected_few_taxa_2
for FILENAME in *.fa
do
awk -F"|" '/^>/{ taxon[$1]++ } END{for(o in taxon){print o,taxon[o]}}'
$FILENAME > $FILENAME\.taxon_count #Creates temporary file with taxon
abbreviation and number of sequences for that taxon in $FILENAME
taxon_count=`grep -v 0 $FILENAME\.taxon_count | wc -l` #Counts the
number of lines with an integer >0 (= the number of taxa with at least
1 sequence)
if [ "$taxon_count" -lt "$MIN_TAXA" ] ; then
echo $FILENAME
mv $FILENAME ./rejected_few_taxa_2/
fi
done
rm -rf *.fa.taxon_count
echo Done
echo
```

```
#Makes a tree for each OG with FastTree
echo "Making a tree for each OG using FastTreeMP..."
for FILENAME in *.fa
do
FastTreeMP -slow -gamma $FILENAME > $FILENAME.tre
done
rename 's/.fa.tre/.tre/g' *.fa.tre
echo Done
```

**PhyloPyPruner:**

```
#Runs PhyloPyPruner 0.9.5
echo "Running PhyloPyPruner..."
phylopypruner --threads $CORES --dir . --min-taxa $MIN_TAXA --min-len
$MIN_SEQUENCE_LENGTH --min-support 0.75 --mask pdist --trim-lb 3 --
trim-divergent 0.75 --min-pdist 0.01 --trim-freq-paralogs 3 --prune MI
cd phylopypruner_output
FastTreeMP -slow -gamma supermatrix.fas > FastTree.tre
cd ..
```

**Extracellular Localization Signals:**

Uploaded the set of 31 genes from SilkSlider as a .fa file to SignalP 5.0 WebServer
